# Top-down and bottom-up controls on soil carbon and nitrogen cycling with repeated burning across four ecosystems

**DOI:** 10.1101/565135

**Authors:** Adam F. A. Pellegrini, Sarah E. Hobbie, Peter B. Reich, Ari Jumpponen, E. N. Jack Brookshire, Anthony C. Caprio, Corli Coetsee, Robert B. Jackson

## Abstract

Fires shape the biogeochemistry and functioning of many ecosystems, but fire frequencies are changing across large areas of the globe. Frequent fires can change soil carbon (C) and nitrogen (N) storage through both “top-down” pathways, by altering inputs through shifting plant composition and biomass, and “bottom-up” ones, by altering losses through decomposition and turnover of soil organic matter. However, the relative importance of these different pathways and the degree to which they regulate ecosystem responses to decades of changing fire frequencies is uncertain. Here, we sampled soils and plant communities in four North American and African sites spanning tropical savanna, temperate coniferous savanna, temperate broadleaf savanna, and temperate coniferous forest that each contained multiple plots repeatedly burned for 33-61 years and nearby plots that were protected from fire over the same period. The sites varied markedly in temperature, precipitation, species composition, fire history and soil chemistry; thus they represent a broad test for the generality of fire impacts on biogeochemical cycling. For all four sites, bulk soil C and N by were 25-180% higher in unburned vs. frequently burned plots, with greater soil losses occurring in areas with greater declines in tree cover and biomass inputs into soils. Fire reduced the activity of soil extracellular enzymes that hydrolyze labile C and N from soil organic matter by two- to ten-fold, whereas tree cover was the predominant control on the oxidation of recalcitrant C compounds. C-acquisition enzyme activity tended to decline with decreasing soil N, suggesting that N losses may contribute to limited decomposition, buffering systems against increased losses of soil C with fire. In conclusion, variability in how fire alters soil C and N across ecosystems can be explained partly by fire-driven changes in tree cover and biomass, but the slower turnover of organic matter we observed may offset some of the reduction of C inputs from plants after fire.

## Introduction

Fires burn ∼570 million hectares of land globally each year, altering the storage and cycling of carbon (C) and nutrients and the composition of organisms (Bond-Lamberty et al. 2007, Bowman et al. 2009, van der Werf et al. 2017, Pellegrini et al. 2018). Many ecosystems burn regularly, with fire-return intervals ranging from two years in tropical savannas to several hundred years in boreal forests (Archibald et al. 2013, Andela et al. 2017). However, fire frequencies are changing, both increasing due to climate change and decreasing due to land-use change (Westerling et al. 2006, Dennison et al. 2014, Andela et al. 2017). Increasing fire frequencies can shift ecosystem C and nutrient cycles by repeatedly combusting plant biomass, volatilizing the C and nutrients in organic matter before they are decomposed into soils (e.g., Ojima et al. (1994), Kauffman et al. (1995), Pellegrini et al. (2014), Muqaddas et al. (2015)). Consequently, fire frequency can alter several belowground properties, such as soil C storage, nutrient availability, and decomposition activity (Guinto et al. 2001, Reich et al. 2001, Hart et al. 2005, DeLuca and Sala 2006, Turner et al. 2007, Jackson et al. 2017). Recent work has demonstrated that repeated burning generally depletes bulk soil C and N but has also helped explain why soil C and N are depleted in some ecosystems, but not others (Pellegrini et al. 2018).

The responses of soil C and N stocks to fire are in part regulated by total plant biomass responses, with larger fire-driven reductions in plant biomass expected to lead to larger reductions in soil C and N (Pellegrini et al. 2014, Kowaljow et al. 2018). Within fire-influenced landscapes, heterogeneity in soil C and N can result from the local enrichment under individual trees, which can have higher soil C and N concentrations and faster decomposition than areas away from tree canopies (Belsky et al. 1989, Ludwig et al. 2004, Dijkstra et al. 2006). Reduced tree cover due to repeated burning (Higgins et al. 2007, Peterson et al. 2007, Holdo et al. 2009a) can reduce the area of soils enriched, thereby lowering soil C and N at the landscape scale (e.g., (Coetsee et al. 2010, Holdo et al. 2011)).

In addition to changing the total amount of C and N entering soils, fire can alter the turnover of C and N within and potential losses from soils through direct combustion and shifting enzyme activity (Treseder et al. 2004, Holden et al. 2013). One hypothesis is that fire accelerates soil C and N turnover by stimulating decomposition activity, potentially through pulsed availability of organic matter inputs after fire (Boerner et al. 2006, Rietl and Jackson 2012). Alternatively, fire may reduce decomposition by lowering microbial biomass through heat-induced mortality, substrate removal, or changes in the microbial community (Waldrop and Harden 2008). In turn, the response of soil organic matter decomposition can be an important factor regulating fire effects on soil C, with suppressed decomposition offsetting the lower inputs of plant biomass (Treseder et al. 2004, Holden et al. 2013).

Yet changes in the decomposition of organic matter likely depend on several factors. For one, the composition of organic matter can be important: fire can enrich lignin-derived but deplete polysaccharide-derived organic matter (Neff et al. 2005), requiring an upregulation of oxidative enzymes (degrading lignin and polyphenols) but not necessarily hydrolytic enzymes (degrading starch, hemicellulose, cellulose) to sustain decomposition. Furthermore, lower soil N availability in burned areas may result in lower organic matter decomposition (e.g., leaf litter (Melillo et al. 1982)). Measuring the responses of extracellular enzymes provides insight into these different factors because it disentangles the potential decomposition of different forms of organic matter and the acquisition of C and N compounds separately.

Here we evaluate the extent to which fire effects on soil C and N are driven by losses in plant biomass inputs compared with changes in decomposition activity. Specifically, we investigated (i) whether landscapes burned repeatedly have lower total soil C and N than those protected from fire, including the degree to which this potential change is caused by direct combustion compared with changes in tree cover and local enrichment (Pathway #1 in Figure 1); and (ii) how fire changes soil C and N turnover, including decomposition of particular forms of organic matter (Pathway #2 in Figure 1). To do so, we sampled plots in four fire-prone ecosystems: tropical savanna, with an overstory composition of broadleaf trees, and two temperate savannas with an overstory composition of either broadleaf or coniferous trees, and a temperate coniferous forest. In each site, we compared plots that had been burned repeatedly with frequencies ranging from decadal to annual for 33-61 years with nearby unburned plots over the same period. The sites varied markedly in temperature, precipitation, species composition, and fire history; presenting a test of the generality of fire impacts on biogeochemical cycling.

**Figure 1:**
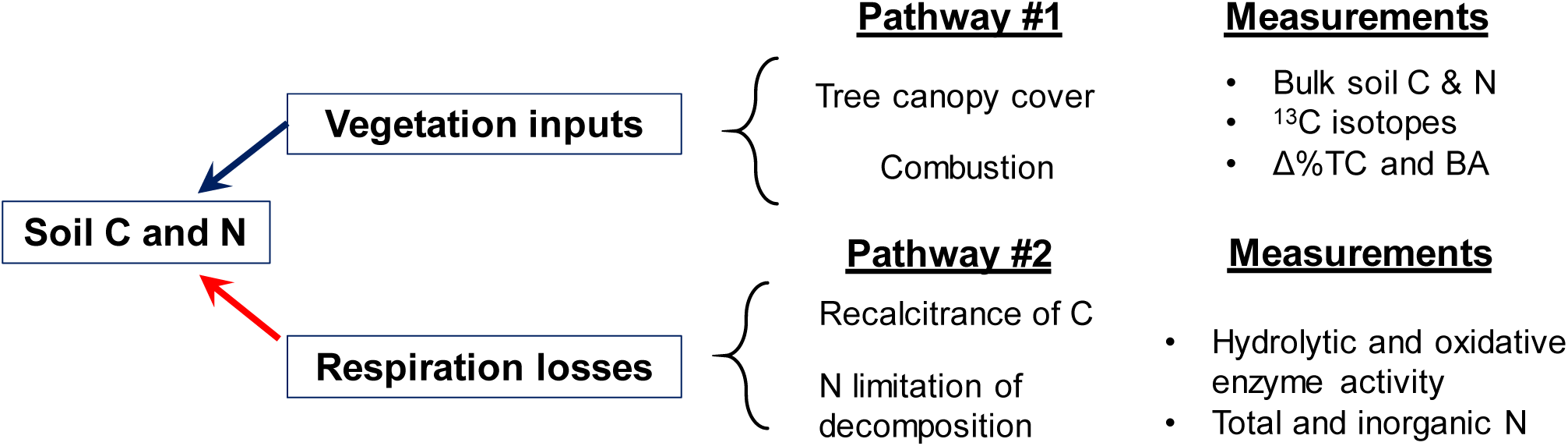
Conceptual schematic of the pathways by which fire may alter soil carbon (C) and nitrogen (N) (denoted Pathway 1 and 2) distinguishing changes in vegetation inputs from respiration losses. To assess the importance of pathway #1, the ‘top-down’ pathway, whereby fire alters the inputs of living and senesced plant biomass into soils, we measured the effect of fire underneath tree canopies vs. in the open to quantify the average effect of fire irrespective of vegetation type (i.e., ‘combustive’ effect) vs. fire-driven changes in tree cover by testing whether trees cause local modifications of soil chemistry (i.e., ‘vegetation type’ effect) and fire changes the amount of tree cover. Measurements of these effects include the bulk soil C and N, the isotopic composition of ^13^C (which gives insight into woody biomass inputs), and the woody plant community (changes in tree cover and basal area). To assess the importance of pathway #2, the ‘bottom-up’ pathway, whereby fire alters the turnover of C in soils, we measured enzyme activities involved in the processing of recalcitrant (polyphenols and lignin) and labile (cellulose, hemicellulose, and starch) C forms. We also evaluated potential effect of N limitation on these processes using measurements of bulk soil N, inorganic N, and N acquisition enzymes.

**Figure 2:**
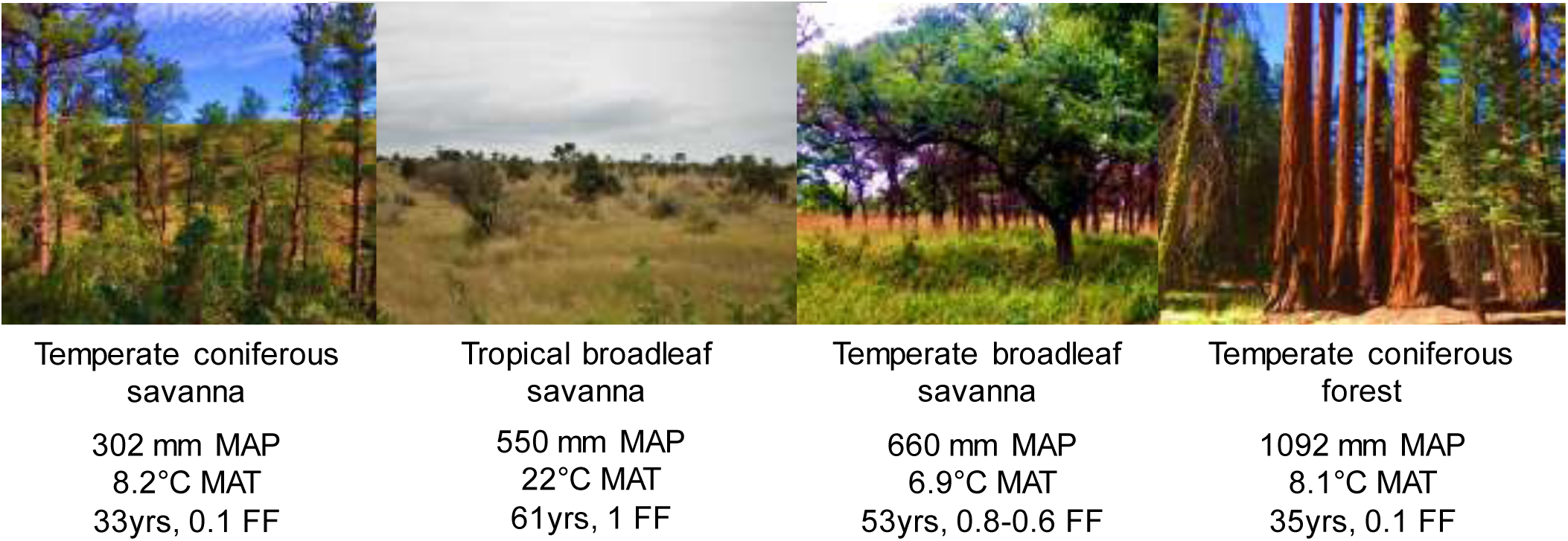
Sites sampled in a temperate coniferous savanna (Missouri Breaks), tropical broadleaf savanna (Kruger), temperate broadleaf savanna (Cedar Creek), and temperate coniferous forest (Sequoia). The sites vary in vegetation structure and climates but have all burned repeatedly at frequencies of 0.1-1 for 33-61 years with adjacent control plots that have not burned over the same period. The values beneath the pictures are the mean annual precipitation (MAP) and temperature (MAT), length of time plots have differed in fire history (years), and the fire frequency of the burned plots (FF).

## Methods

We sampled sites in four locations spanning a range of forests and savannas that experience frequent burning: temperate coniferous forest, temperate coniferous savanna, temperate broadleaf savanna, and tropical broadleaf savanna. In each site, we sampled three replicate plots of each fire treatment (plots protected from fire and those burned repeatedly). In all sites except the temperate coniferous savanna, the replicates were distinct and separated by fire breaks. Within each plot, we sampled both directly underneath tree canopies and away from canopies to test the impact of changing woody plant abundance. Where more than one tree species was locally abundant, we replicated our sampling across different tree species to test for interspecific variability in tree effects on soils, as described below. We sampled three individuals for each tree species when they were sufficiently abundant.

### Site descriptions

The tropical broadleaf savanna site was located in Kruger National Park, South Africa in the ‘Experimental Burn Plots,’ which have received different fire frequency treatments since the mid-1950s (1956 at the particular site we sampled in 2017). The large-scale plots are ∼5-7 ha in size and are replicated across the park (Biggs et al. 2003). Plots are exposed to a suite of meso- to mega-fauna. We sampled plots located in the Skukuza area (latitude: −25.10, longitude: 31.45), that receive around 550-600 mm yr^-1^ with a mean annual temperature of ∼22°C. In March of 2017, we sampled sites on plots burned annually at the end of the dry season in August, when fires are most intense, and compared these to plots that have remained unburned since the onset of the experiment. The plots are dominated by broadleaf woody plant species in the genera *Combretum* and *Terminalia*, which are common tree species in savannas of the region. The sites span a large fire-driven gradient in tree biomass and grass cover is substantial across all plots, even in those protected from fire. We sampled soils in grassy areas as well as underneath the canopies of four different tree species: *Combretum apiculatum, Combretum collinum, Combretum hereroense*, and *Terminalia sericea.*

The temperate broadleaf savanna was in the Cedar Creek Ecosystem Science Reserve in Minnesota, USA (latitude: 45.40, longitude: −93.19). The fire plots were established in 1964 on a landscape that was primarily savanna and woodland. Large areas of several hectares were delineated with fire breaks and assigned to different fire frequency treatments ranging from complete exclusion to burning every 8 out of 10 years or so. Plots are burned in the spring (ranging from April-May depending on conditions). This site receives 660 mm yr^-1^ in precipitation and has a mean annual temperature of 6.9°C. The plots are exposed to deer herbivory. The tree communities are dominated by broadleaf species including bur oak and northern pin oak (*Quercus macrocarpa* and *Q. palustris*, respectively). Consequently, the plots we sampled in August of 2017 spanned a fire-driven ecotone between open savanna, with continuous grass cover in the high frequency plots, to closed-canopy forest without a continuous grassy layer under trees in the unburned plots (Peterson and Reich 2001). We sampled underneath tree species *Q. macrocarpa* and *Q. palustris* and in grassy areas outside of tree canopies. In the unburned plots where forests have formed, we sampled in areas not yet encroached by forest.

The temperate coniferous savanna was in the Missouri Breaks region in central Montana, USA. These plots do not receive direct fire manipulation, so we used variability in wildfire history to examine different fire regimes. Using remote-sensing records dating from 1984 [Monitoring Trends in Burn Severity product, which has been used to establish trends in wildfire, (Dennison et al. 2014)] we identified sites that have similar underlying geologies and climates (latitude: 47.43, longitude: −108.17). The burned plots were established inside perimeters of fire scars that burned each decade since 1984, with the last fire in 2012. We determined the fire history by layering fire scars and sampling with the areas that had repeatedly burned. In the unburned plots, we know that they had not burned since at least 1984, but have no data on the exact date of the last fire. Plots typically burn in the late summer. This site receives 302 mm yr^-1^ in precipitation and has a mean annual temperature of 8.2° C. The landscape contains bare areas, rock outcrops, and shrublands, which we avoided in our sampling. We sampled underneath the main woody plant species, *Pinus ponderosa* and *Juniperus scopulorum*. In the unburned plots, thick litter layers can accumulate underneath trees, excluding understory grasses, which were largely absent in the burned plots; however, the ‘open’ areas we sampled were always positioned in the grassy matrix. These sites were sampled in June 2017. The large area with different fire histories (distributed across a 15 ha area for the unburned ‘treatment’ and across a separate 10 ha area for the burned ‘treatment’ that identified to be within fire scars that delineated repeatedly burned areas) allowed us to segment plots within the landscapes containing different fire frequencies, established in plots 100-400 m apart from one another.

The temperate coniferous forest was in Sequoia and Kings Canyon National Parks in California, USA. Fire effects plots, which have experienced different historical fire regimes (e.g., (Schwilk and Caprio 2011)), were established in the national parks, with surface fuels, vegetation, and fire behavior monitored pre- and postfire. Plots are 10 ha in size and were established in the 1980s to include a variety of vegetation types. We sampled plots where fire has been excluded for >150 years (Caprio and Swetnam 1995, Swetnam et al. 2009) and where prescribed burns have been conducted every 10-15 years (plots located in Giant Forest, latitude: 36.60, longitude: −118.73), starting in 1983-1986 for these sites. We sampled plots in a forested landscape currently dominated by *Abies concolor* and *Sequoiadendron giganteum*. There is little understory vegetation in plots where fire has been excluded, which have deep accumulations of litter, duff, and coarse wood debris, but understory plant species such as *Ceanothus, Lupinus*, and various grasses occur more frequently in plots where fire has been restored. Plots have usually been burned by prescribed fire during either the spring or fall (May-June or September-October). The study area has a Mediterranean climate, with the sites sampled at ∼2100 m elevation receiving on average 1092 mm in precipitation (1920-2017 at Giant Forest USACE station). Most precipitation occurs during the winter (∼50% as snow) with dry summers and a distinct period of water deficit (Stephenson 1988). Average annual air temperatures range from 1.9-13.6°C in Grant Grove at a similar elevation to Giant Forest (van Mantgem et al. 2016). These sites were sampled in September 2017. We sampled underneath *Abies concolor* and *Sequoiadendron giganteum* as well as in ‘open’ areas. The open areas were outside of tree canopies and dominated by herbaceous and shrubby species in the burned plots; however, in the unburned plots the samples were away from tree trunks but still underneath thick litter and duff layers because of the lack of fire for >150 years.

### Soil sampling

At each sampling location within a site (e.g., under an individual of a giant sequoia), we aggregated five soil cores from the top 5 cm of the mineral horizon sampled in the shape of a 125cm^3^ cube (5×5×5cm for each sample, total volume of 625 cm^3^ per-sampling location). We identified the mineral horizon by first digging a hole >20 cm in depth and removing the organic horizon from the profile, when present, and sampling the top 0-5 cm of the mineral horizon. We focused on the upper soil horizons because they have the potential to be most responsive to fire and the most biologically active.

Soils were passed through a 2 mm sieve to remove coarse particles and partitioned into three sub-samples: (i) fresh soil for soil moisture and inorganic N analyses, (ii) frozen soil for enzyme analyses (see below), and (iii) dried soil for total C and N and ^13^C analyses. Moisture was determined by drying soils at 105° C until they reached a constant weight (24-48 hours). We used values of moisture content to adjust all analytical chemistry values to grams per-dry weight of soil.

### Vegetation sampling

We analyzed the response of the woody plant communities in the tropical and temperate broadleaf savannas as well as the temperate coniferous forest sites. In these three sites, tree cover had a significant effect on several soil variables (see *Results*) and thus changes in woody cover were important for explaining shifts in soils at the landscape scale. Unfortunately, we do not have any surveys on the woody plant community in the temperate coniferous savanna; however, we did not observe any effect of tree cover on soil C, N, or enzyme activities. In the tropical savanna, we ran two belt transects that covered 300 m^2^ each and measured the height and diameter at 30cm of every individual tree (greater than 30 cm in height). The crown area was calculated for these trees using allometric equations for species within the park.

In the temperate broadleaf savanna, the woody plant community is surveyed by the Cedar Creek Long Term Ecological Research program every five years within 3,750 m^2^ permanent plots established in the replicate plots for each fire treatment in 1995. On each survey, stem diameter was measured at ∼1.35 m. We used the measurements from the most recent survey in 2015. We also compiled data from previous measurements of canopy cover determined by measurements of light penetration using a pair of LICOR LAI-2000 sensors. To measure the effects of trees alone, the sensor was placed above the tall shrub layer (usually 0.5-1 m in height). Readings were taken under conditions of uniform, diffuse light (twilight hours, or on cloudy days) and are paired with readings from an “above” sensor operating continuously (one reading every 15 seconds) in a nearby open field.

In the temperate coniferous forest, we also used data collected by the National Park Service following Fire Monitoring Handbook protocols (USDI National Park Service 2003) on tree basal area, litter, and duff (partially decomposed litter above the soil organic horizon) to evaluate whether the lack of a response in soil properties was associated with no change in tree basal area. Each permanent monitoring plot is located directly adjacent (within 100 m) to the corresponding plot we sampled for soils; we sampled soils outside of the permanent plots to avoid disturbance. Tree communities were surveyed inside each replicate plot, with the first survey in the burned plots taking place the year before the first burn and subsequent measurements immediately after the burn and again at 1-, 5-, and 10-year intervals. In the unburned plots, surveys started in 2007, and we used the survey data from 2012. Given the important contributions of litter and “duff” (partially decomposed litter above the soil) layers to ecosystem stocks in these forests, we also analyzed duff and litter biomass measurements taken in the same plots and at the same time as when the tree communities were surveyed.

### Analyses of C and N in soil and C isotopes

Total soil C, N, and ^13^C were measured by combusting the sample using a Costech coupled Elemental Analyzer and Mass Spectrophotometer at Stanford University and University of Maryland. Mass combusted per-sample was determined to maximize analytical accuracy of measurements. Duplicates were run for 10% of all samples to ensure analytical precision which we defined to be an error of less than 5%.

We measured inorganic N (IN) on soils within 48 hours of collection. Inorganic N was measured by extracting ∼5 g of freshly sampled soil via shaking in 50 ml of 1M KCl. Following shaking, the samples were centrifuged and filtered through Grade 41 ashless Whatman filters. Inorganic N was analyzed on an automated spectrophotometer WestCo SmartChem 200 discrete analyzer at Stanford University. Nitrate was analyzed using a cadmium column reduction, followed by a diazotization with sulfanilamide coupled with N-(1-naphthyl)ethylenediamine dihydrochloride, which was analyzed colorimetrically at 550 nm. Ammonium was analyzed using the indophenol blue method, where it reacts with salicylate and hypochlorite in the presence of sodium nitroferricyanide to form the salicylic acid analog of indophenol blue in a buffered alkaline solution (pH 12.8-13), which was analyzed colorimetrically at 660 nm.

### Enzyme activity

To assess potential turnover of C and N, we measured hydrolytic and oxidative extracellular enzyme activity in a subset of samples at each site. The hydrolytic enzymes we measured were: cellobiohydrolase (CBH, degrades cellulose), β-glucosidase (BG, degrades cellulose), α-glucosidase (AG, degrades starch), β-xylosidase (BX, degrades hemicellulose), and N-acetyl-β-D-glucosaminidase (NAG, degrades chitin). The oxidative enzymes were: phenol oxidase (PO, degrades polyphenols), and peroxidase (PX, degrades polyphenols/lignin). The enzyme analyses were performed using methods presented in Hobbie et al. (2012), which used slightly modified methods of Sinsabaugh et al. (1992) and Saiya-Cork et al. (2002), and were performed at the University of Minnesota.

Enzyme activities were analyzed both individually and by taking the sum of all hydrolytic C acquisition enzymes and the oxidative C acquisition enzymes separately. We refer to the group of hydrolytic enzymes as degrading ‘labile’ substrates and the oxidative enzymes as degrading ‘recalcitrant’ substrates because of the higher energy required to breakdown the polyphenols and lignin targeted by oxidative enzymes (similar to Sinsabaugh et al. 2008). We also analyzed the ratios of enzyme activities to ascertain how fire and vegetation type may be altering the relative turnover of C vs. N and the hydrolytic C vs. oxidative C enzymes. These ratios illustrate which types of enzymes are more sensitive to change and shifts in the cycling of C and N (Sinsabaugh and Follstad Shah 2011).

### Respiration

To link measurements of enzymes with soil C losses, we took advantage of previous measurements on soil C respiration across the temperate broadleaf savanna experimental plots at Cedar Creek conducted from 1999-2005. Because the respiration measurements did not change directionally with time, we assume that the variability across plots from the 1999-2005 are representative of present-day conditions. These measurements were made throughout the growing season (April-October) across eight points within each replicate plot. Aboveground biomass was removed above each point prior to measurement, which were made using a Li-Cor 6400-09 soil respiration chamber attached to a Li-Cor 6200 gas exchange system over a 5 cm permanent soil collar. We re-scaled the data to fluxes per-degree Celsius because of the high correlation between respiration and temperature (F_1,222_=1703, p<0.0001).

### Data analysis

To test the effect of fire and vegetation type on soil chemistry, we used mixed effects models to take advantage of the hierarchical sampling design of individuals within species within plots across fire treatments (Bates et al. 2014) (package *lme4*). Because of the stratified sampling of several individuals within a single replicate plot of a treatment within a site, we included replicate plot as a random intercept. The fixed effects of fire, vegetation, and their interactions were determined using model selection via AIC with a threshold of two. When AIC values did not differ by more than 2 and the lower AIC model was more complex, we used a χ^2^ test to determine whether the more complex model added significant explanatory power. If the test was insignificant, we used the simpler model. Significance of terms was evaluated using the package *lmerTest* that approximates the degrees of freedom using Satterthwaite’s method (Kuznetsova et al. 2017). Overall tests across all sites also included a ‘site’ effect as an additional fixed effects variable in the model selection process to test whether sites differed in soil chemistry and sites had different responses to either fire or vegetation type (i.e., site-by-fire interaction). The data were log-transformed when necessary to fulfill assumptions that residuals were normally distributed.

To quantify the relative effects of changes in canopy area, we weighted soil concentration data by tree canopy cover to calculate landscape-level averages of soil C and N in the different fire treatments. For example: [N]_weighted_=[N]_tree_*tree cover+[N]_open_(1-tree cover), where tree cover is a proportion of the total landscape area (ranging from 0 to 1). This weighted value was compared to the average values across soils underneath grasses and trees (i.e., not weighting samples by canopy cover).

We use ^13^C to infer contributions of tree biomass to soils, which assumes that most grasses are C4. In many savanna ecosystems, the strong fractionation differences between trees (C_3_ photosynthetic pathway) and grasses (mostly the C_4_ photosynthetic pathway), makes ^13^C a useful tracer for understanding how fire-driven reductions in tree biomass lead to losses of soil C. The geographic domain of this approach is limited in its utility in northern latitudes dominated by C_3_ graminoids and thus this would underestimate grass contribution to soil C in the temperate coniferous savanna especially; however, in the tropical and temperate broadleaf savannas ^13^C is a good indicator because C_4_ grasses dominate grass biomass. While pyrolysis of organic matter can also result in ^13^C fractionation (Bird and Ascough 2012), these effects are thought to be relatively small compared to fractionation via the C_4_ vs. C_3_ photosynthetic pathway.

We refer to the vegetation type effect as the contrast between soils taken from underneath tree canopies vs. outside of the canopy. We refer to the fire or combustive effect as the contrast between soils in the burned vs. unburned plots underneath both grasses and trees.

## Results

### Changes in soil C and N driven by both combustion and shifts in tree cover

Across all sites, fire and vegetation type were both important predictors of bulk soil C (p=0.029 and p<0.0001) and N (p<0.0001 and p=0.0275) concentrations; fire tended to deplete soil C and N, and trees tended to enrich C and N when all sites were analyzed in a mixed-effects model. When each site was analyzed independently for fire and vegetation type effects, combustion effects alone (defined to be the effect of fire on soils within vegetation types) did not always have a clear effect on soil chemistry; rather, soil properties in certain ecosystems were driven primarily by changes in tree canopy area because of the localized effects of trees on soils, which tended to become increasingly important in the wetter ecosystems.

In the site with the lowest precipitation, the temperate coniferous savanna at Missouri Breaks, changes in soils were primarily caused by combustive effects independent of vegetation type (Tables 1,2, Figure 3). Fire exclusion plots had 115% higher soil C concentrations than burned plots, but there was only a marginally significant (p=0.08) enrichment effect of soil C underneath trees (+29%) (Tables 1,2, Figure 3). Nitrogen responded similarly, with 178% more soil N in unburned than burned plots and no difference between vegetation types (Tables 1,2, Figure 3).

**Table 1:** Summary of significant responses to either fire, vegetation type, or both for all response variables analyzed. Percentage change is indicated by % values for fire (denoted by the F) for samples from the unburned plot relative to the burned plot and for vegetation (denoted by the V) samples from trees relative to away from the canopy; consequently, positive values indicate enrichment in either unburned plots and underneath trees. The “mix” indicates when the interaction between fire and vegetation type altered the sign of the effect (e.g., enrichment underneath trees in unburned plots, but depletion in burned plots). “T.” or “Tr.” indicate temperate or tropical, respectively, and “conif.” vs. “broad.” indicate coniferous vs. broadleaf tree types. Vegetation-type effect in tropical savanna, p=0.08.

| Response variable |  | T. conif. savanna | Tr. broad. savanna | T. broad. savanna | T. conif. forest |
| --- | --- | --- | --- | --- | --- |
| Total concentrations | Total %C | F+115% | F+29% V+89% | V+99% | n.s. |
|  | Total %N | F+178 | F+38% V+59% | V+62% | V-16% |
|  | C:N |  | V+14% | V+20% | V+11% |
| | $^{13}\text{C}$ | F -49%<br>V+7% | F-8% V-23% | F-8% V-15% | F-3% V-2% |
| Enzyme activity | $\text{C}_{\text{hydro}}$ | F+552 | n.s. | V+68% | F+81% |
| | $\text{C}_{\text{oxi}}$ | n.s. | n.s. | V+175% | V+134% |
|  | NAG | F+941% | F+101% | V+138% | F+192% V+182% |
| N-acquiring/<br>C-acquiring enzymes | NAG/$\text{C}_{\text{hydro}}$ | n.s. | n.s. | V+32% | F+61% V+17% |
| | NAG/$\text{C}_{\text{oxi}}$ | F+480% | V+40% | Mix | F+230% |

**Table 2:**
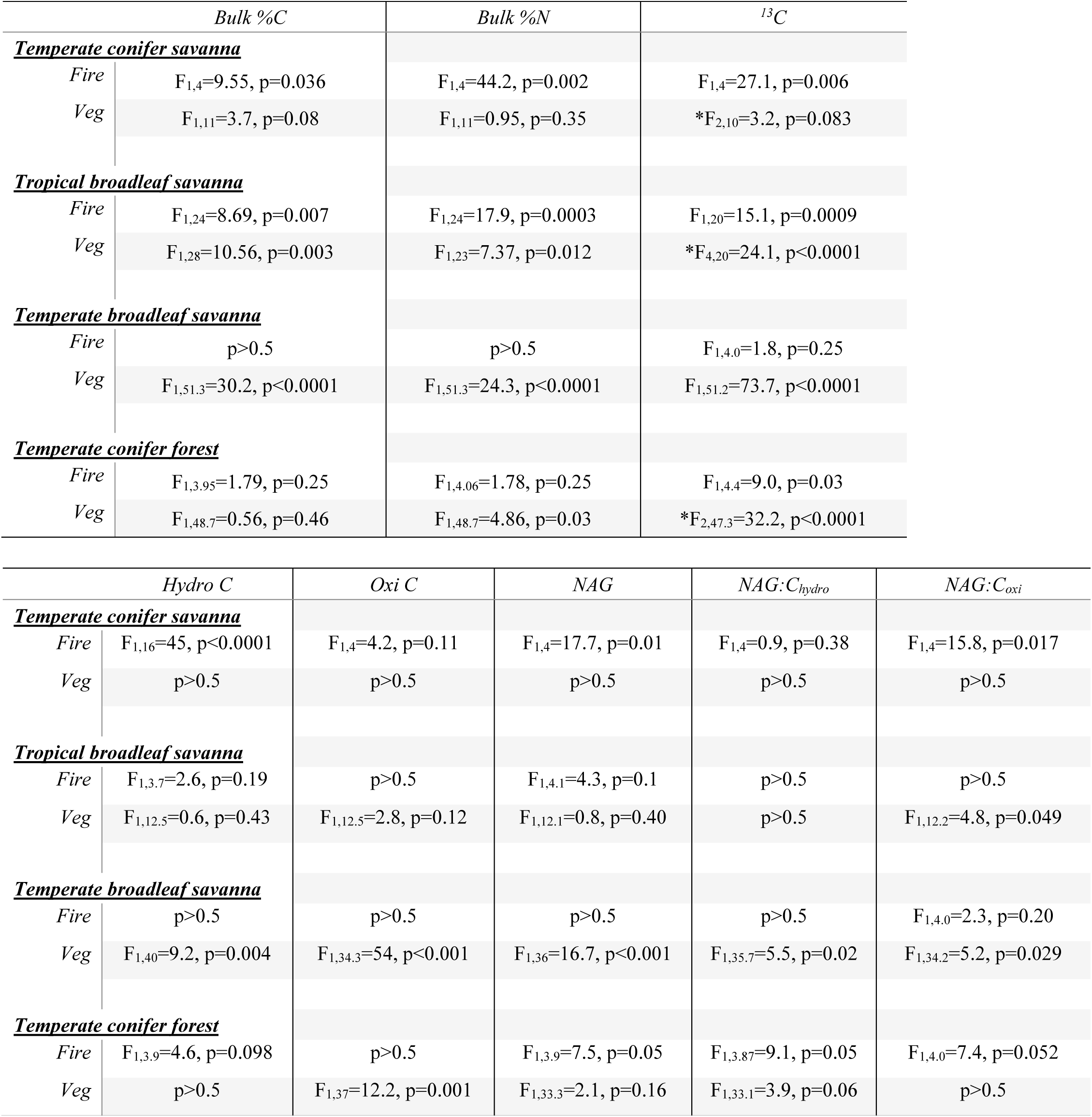
Statistical results from mixed-effects models on soil chemistry variables. “Fire” and “Veg” refer to the comparisons of burned and unburned sites vs. grasses and trees, respectively. Interaction terms, when significant, are presented in the main text. Enzyme variables: sum of hydrolytic C acquisition enzymes (hydro C), sum of oxidative enzymes (Oxi C), enzyme that can acquire N (NAG), ratio between hydrolytic C acquisition to NAG (NAG:C_hydro_), ratio between oxidative C acquisition to NAG (NAG:_Coxi_).

**Figure 3:**
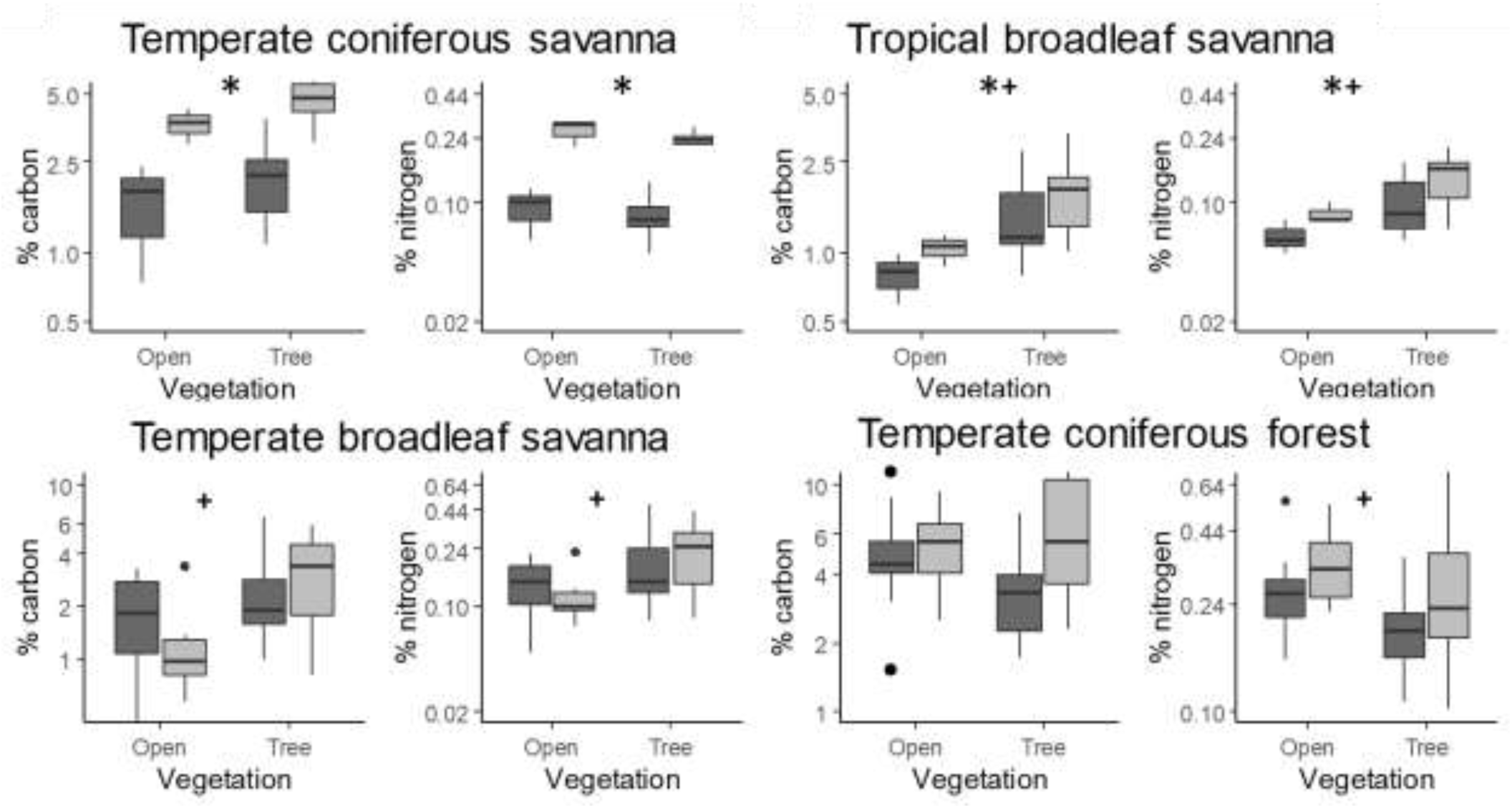
Differential role of fire and vegetation type on bulk soil carbon and nitrogen concentrations across ecosystems. Box plots of soil total carbon and nitrogen to 5 cm depth partitioned into different fire treatments (dark grey=burned, light grey=unburned) from either underneath trees or beyond tree canopies. Y-axes are on a log scale. Significant effect of fire denoted by the asterisk, significant effect of vegetation type denoted by the plus symbol, dots above box plots indicate outliers.

The tropical broadleaf savanna at Kruger National Park, which receives more precipitation than the temperate coniferous savanna but has ∼2.7-fold higher mean annual temperate experienced combined effects of fire and vegetation: there was 29% higher soil C concentrations in unburned vs. burned plots and 82% higher C concentrations underneath trees than in the open. Similarly, unburned plots had 38% higher soil N concentrations than burned plots and 59% more N under trees vs. in the open (Tables 1,2, Figure 3). Consequently, in tropical savannas there are combined effects of both fire and vegetation cover on soil C and N (we quantify the landscape-scale effect of changing tree cover below).

The temperate broadleaf savanna at Cedar Creek, which receives similar rainfall to the tropical savanna but has three-fold lower mean annual temperature, responded to effects of vegetation type in ways that interacted with fire treatments. Soils underneath trees were enriched in C and N by 99% and 52%, respectively, relative to away from tree canopies (Tables 1,2, Figure 3). Although there was not an effect of fire on soils directly (Table 2), fire did modify the enrichment effect of vegetation, with the elevation of soil C underneath trees being greater in unburned plots than in burned plots (interaction term: F_1,50.1_=5.53, p=0.02).

In the temperate coniferous forest in Sequoia National Park, which receives the highest amount of precipitation, total soil C and N tended to be resistant to effects of fire and vegetation. First, there were no significant effects of either fire or vegetation on total soil C (Tables 1,2). Second, although the vegetation type significantly affected soil N, the effects were in the opposite direction: soils in open areas had 21% higher N concentrations than under trees. Fire had no effect on total soil N (Tables 1,2, Figure 3).

The comparison across all the sites illustrated a gradient of responses, with sites ranging from being primarily determined via combustion effects alone in the temperate coniferous savanna, to both combined combustion and tree cover effects in the mesic tropical savanna, to tree cover effects alone in the wet temperate broadleaf savanna, to little change in the temperate coniferous forest.

### Belowground changes correspond with changes in tree populations sizes

Given that fire alters the biomass of vegetation and tree cover, we next wanted to determine the effect of variability in vegetation structure and composition on belowground changes within ecosystems (e.g., how does the local enrichment effect underneath trees scale to landscape-level changes). The first component tests how fire-driven changes in soil chemistry at the landscape scale depend on the differences in soil chemistries underneath trees or in the open observed in the tropical and temperate broadleaf savannas. The second component tests whether the lack of an effect of fire on soil C and N in the temperate coniferous forest may be due to lack of change in vegetation biomass stocks.

In both the temperate and tropical broadleaf savannas, fire strongly reduced tree basal area, abundance, and tree cover. In the tropical savanna, the unburned plots had 60-fold higher tree basal area, 2.9-fold more individuals, and 61-fold higher tree canopy cover than in the burned plots. In the temperate broadleaf savanna, changes in tree populations were also significant: unburned plots had 1.4-fold higher tree basal area, 4.9-fold more individuals and 0.8-fold higher tree canopy cover than burned plots (Tables S2,3, Figure 4).

**Figure 4:**
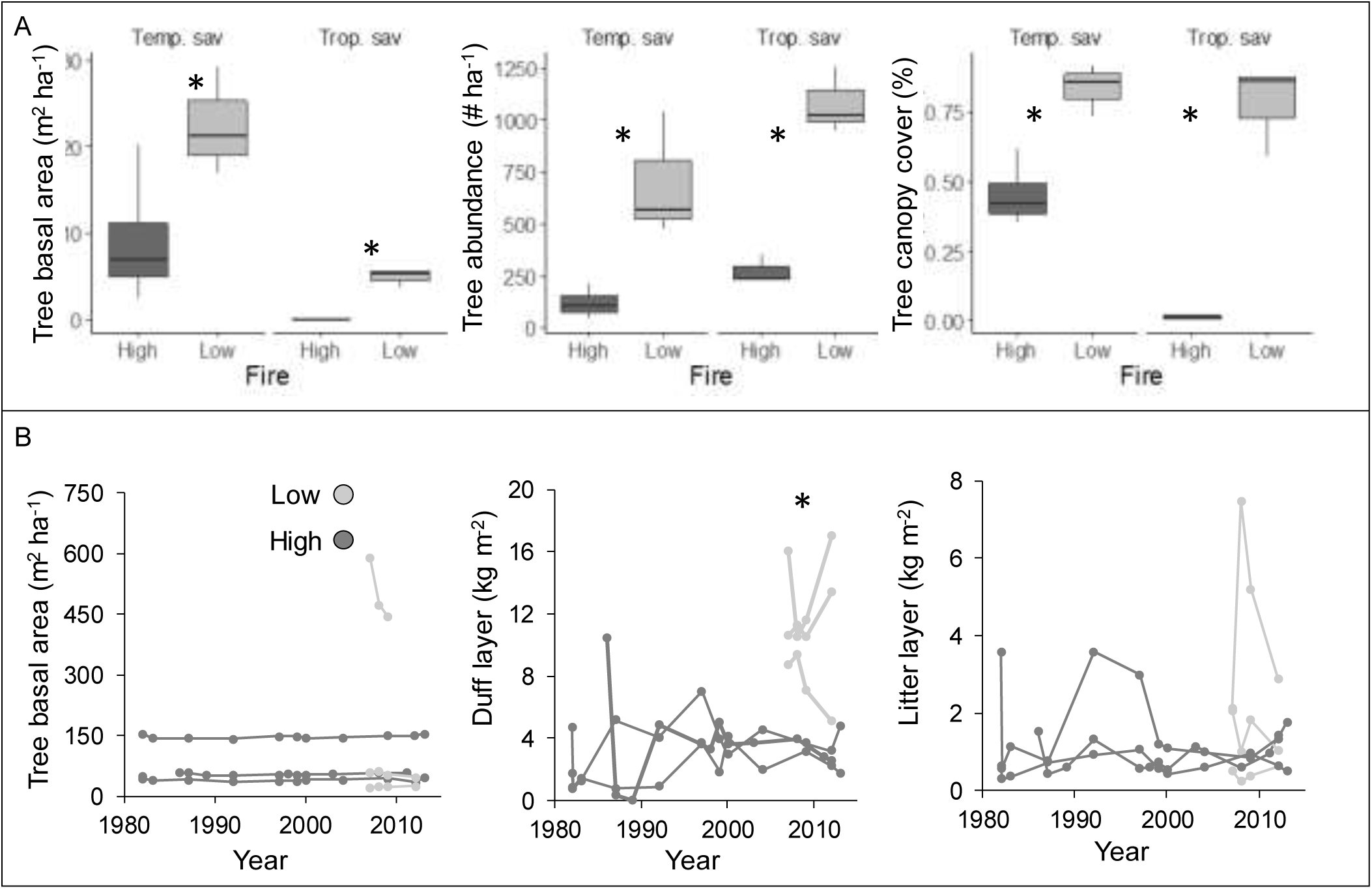
Fire reduces tree basal area, abundance, and structure in tropical and temperate broadleaf savannas, while the temperate coniferous forest tree basal area and biomass are more resistant to fire. Panel A: Tree basal area, abundance, and canopy cover in the different fire treatments in broadleaf temperate and tropical savannas using box and whiskers plots. Panel B: Trends in tree basal area, litter, and duff layer in the coniferous forest, where unburned plot measurements were only conducted since 2007. Dark grey indicates burned and light grey indicates unburned treatments; asterisks denote significant differences between treatments (the temperate broadleaf savanna basal area comparison p=0.08). Lines do not denote significant trends.

Fire-driven declines in tree cover amplify declines in soil C and N in the tropical and temperate broadleaf savannas. In the tropical and temperate broadleaf savannas, soils under trees had higher C and N concentrations than soils in grassy areas suggesting changes in tree cover are important for differences in soil C and N at the landscape scale. When we weighted soil C and N concentrations with tree cover ([value underneath tree]*[proportion tree cover] + [value outside of tree canopy]*[proportion open cover]), we found soil C and N concentrations were 110% and 100% higher in the unburned than in burned plots in the tropical broadleaf savanna. Similarly, in the temperate broadleaf savanna, when we accounted for the higher tree cover in unburned plots and the enrichment of soil C and N underneath trees, the unburned plots had 28% and 25% more C and N than burned plots. The change observed in the temperate broadleaf savanna arose because estimates of soil C and N concentrations weighted by tree cover were 31% and 23% greater, respectively, in unburned plots than estimates assuming an equal distribution of grasses and trees. Consequently, shifting tree cover is a critical component in driving changes in soil C and N at the landscape scale in relatively mesic savannas.

We next tested whether the lack of changes in soil C and N in the temperate coniferous forest might have resulted from lack of changes in plant biomass inputs. Inventory data on trees revealed that generally the two fire treatments had similar tree basal areas. In two of the six plots, the presence of giant sequoias resulted in exceptionally high basal areas (444.5 m^2^ ha^-1^ in one unburned plot and 154.9 m^2^ ha^-1^ in one burned plot, Figure 4). In the other four plots, burned plots had basal areas from 44.6-45.1 m^2^ ha^-1^ and unburned plots had basal areas from 24.8-46.2 m^2^ ha^-1^ (Figure 4). Furthermore, analyses of basal area trends through time illustrated that fire treatment did not affect the change in basal area through time (F_2,40.6_=1.3, p=0.29, Figure 4). Consequently, there was little evidence that fire reduced tree basal area.

Within the temperate coniferous forest, we also tested for differences in the mass of litter and duff (partially decomposed litter that has accumulated on the soil surface) layers, which can be substantial pools of C and N (Figure 4). Unburned plots had ∼4-fold higher duff stocks than the burned plots (10.9 vs. 2.9 kg m^-2^, F_1,4.6_=15.7, p=0.013). However, unlike the duff layers, fires had no significant effects on litter stocks (F_1,4.6_=0.59, p=0.48).

Taken together, comparisons across the sites reveal that dynamics of tree biomass inputs are important for explaining the variable responses in soil C and N responses to fire. In the mesic savannas, shifts in tree cover amplified fire effects on soils at the landscape scale, and in the temperate coniferous forest the stability of tree populations and litter layers is a possible mechanism leading to no change in soils with fire.

### Losses of soil carbon correspond with lower tree biomass inputs

We next utilized ^13^C to further test how changes in soil C depended on tree biomass inputs into soils in the savannas where the different fractionations between trees and grassed allowed us to partition the contribution of tree biomass to soil C. The relationship between losses of soil C and lower inputs of tree biomass into soils was supported by correlations between ^13^C and total C. In all sites there was a significant negative correlation between soil C and ^13^C (temperate broadleaf savanna: F_1,51.2_=40.0, p<0.0001; temperate conifer savanna: F_1,13.9_=6.8, p=0.021; tropical savanna: F_1,28_=5.3, p=0.029). Because trees are depleted in ^13^C relative to grasses, the inverse correlation suggests a link between lower C and a decline in tree biomass inputs. Furthermore, analyses of fire and vegetation type effects within each site illustrated that soils from unburned plots and underneath trees had lower ^13^C (Tables 1&2). Taken together, the fire-driven reductions in tree cover and biomass inputs into soil organic matter were important factors explaining the variability in fire effects on soil C and N across sites.

### Fire reduces extracellular enzyme activity

We hypothesized that the activity of extracellular enzymes involved in breaking down organic matter would be reduced in burned plots and away from tree canopies and would respond to fire and vegetation type in similar ways as bulk soil C and N responded. When all sites were considered, the cumulative activities of enzymes that break down high energy-yielding compounds such as cellulose, hemicellulose and starch were lower in burned than in unburned plots (p<0.0001).

In the temperate coniferous savanna, total C acquisition enzyme activity was 550% higher in the unburned plots than the burn plots (Tables 1,2, Figure 5), similar to the response of total soil C. In the temperate broadleaf savanna, hydrolytic C acquisition activity was 68% higher underneath trees than in the open (Tables 1,2, Figure 5); combining the effect of trees with changes in tree cover in the temperate broadleaf savanna, we estimated that unburned plots had 49% higher activities at the landscape scale. Similarly, in the temperate coniferous forest, total hydrolytic-enzyme activity was 81% higher overall in unburned plots, with the effect significantly different in soils under trees (Figure 5; fire-vegetation interaction: F_1,32.1_=9.2, p=0.005). Only in the tropical savanna, total hydrolytic enzyme activity was not affected by fire or vegetation type (Tables 1,2, Figure 5). However, unchanged total activity masked the variation in the responses of the individual enzymes: CBH and BX, which degrade cellulose and hemicellulose, respectively, tended to be lower in burned plots in the tropical savanna. Taken together, fire reduced total activity of C-degrading hydrolytic enzymes at the landscape-scale in three of the four sites, but always reduced at least two or more individual hydrolytic enzymes at every site.

**Figure 5:**
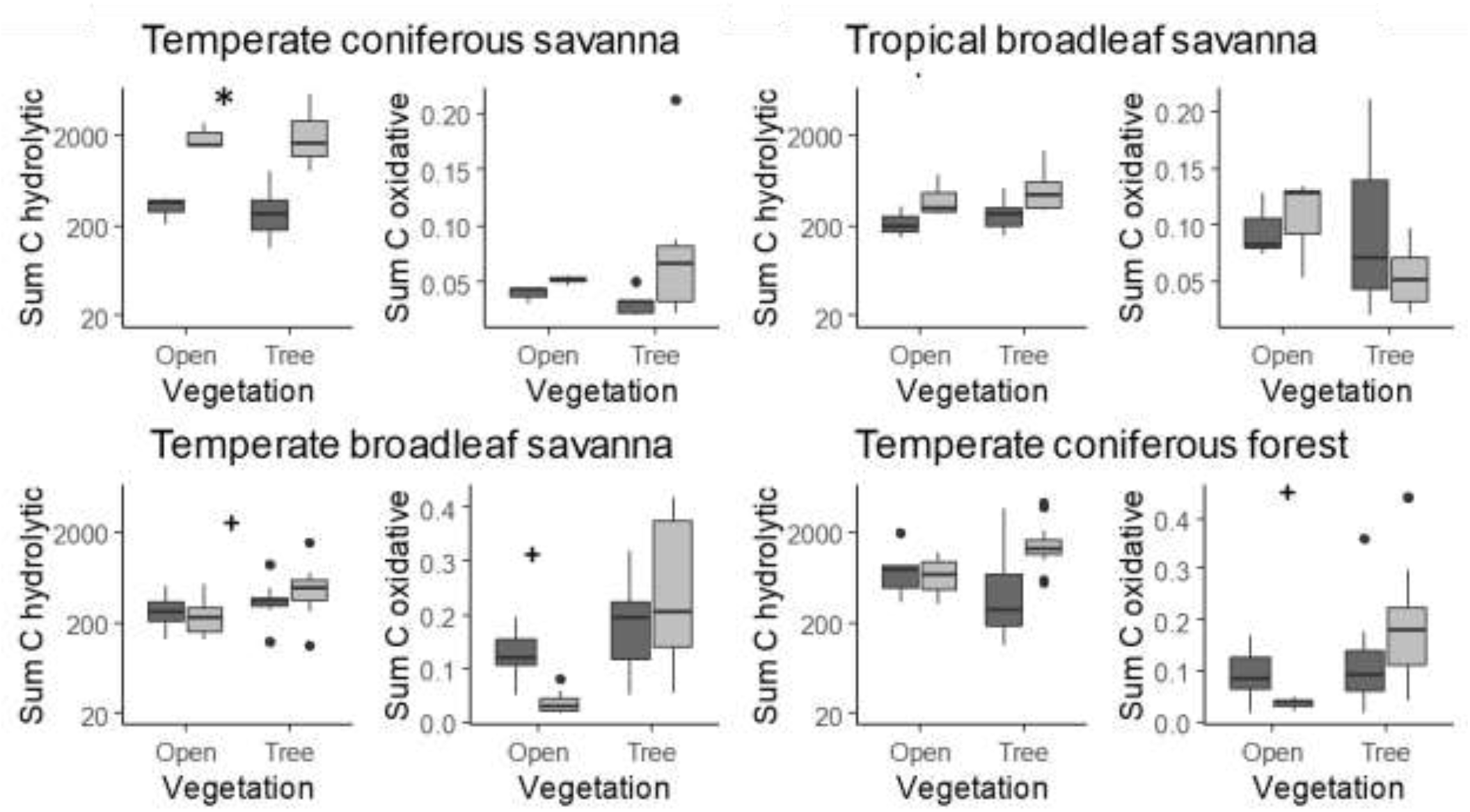
Fire and tree cover can influence activity of enzymes that degrade labile C compounds, whereas tree cover influences enzymes that decompose recalcitrant C compounds. Box and whiskers plots of six extracellular enzymes aggregated into hydrolytic and oxidative categories. The hydrolytic enzymes are: cellobiohydrolase (degrades cellulose), β-glucosidase (degrades cellulose), α-glucosidase (degrades starch), and β-xylosidase (degrades hemicellulose). The oxidative enzymes are: phenol oxidase (degrades polyphenols), and peroxidase (degrades polyphenols/lignin). Values partitioned into different fire treatments (dark grey=burned, light grey=unburned) from either underneath trees or away from tree canopies. Significant effect of fire denoted by the asterisk, significant effect of vegetation denoted by the plus symbol. Units for hydrolytics are nmoles h^-1^ g^-1^ and units for oxidatives are μmoles h^-1^ g^-1^.

In contrast to C-degrading hydrolytic enzymes, the activities of oxidative enzymes, which degrade more recalcitrant C compounds, were resistant to fire but responded to vegetation type. For example, oxidative enzyme activities were 134% and 175% higher under trees than in open areas in the temperate coniferous forest and temperate broadleaf savanna, respectively (Tables 1,2, Figure 5). Because fire reduced tree cover in the temperate broadleaf savanna, oxidative enzyme activity was 40% higher at the landscape scale in the unburned plots because of higher tree cover. In the coniferous savanna and tropical savanna, neither vegetation type nor fire affected total oxidative enzyme activity (Tables 1,2, Figure 5).

Although sites differed in how fire and vegetation type affected enzyme activity, changes in extracellular enzyme activity were positively coupled with changes in bulk soil C. There was a positive correlation between hydrolytic enzyme activity and soil C concentrations in each site (Table S5), even when the statistical model incorporated fire or vegetation type effects. The response of oxidative enzyme activity was less strongly related to changes in bulk soil C, with only a significant positive correlation in the temperate broadleaf savanna (Table S5).

The contrasting responses of enzymes (both to fire and vegetation type as well as correlations with bulk soil C) illustrate that hydrolytic and oxidative enzyme activities are controlled by different factors. Hydrolytic enzymes appear to scale with the amount of soil organic matter, whereas oxidative enzymes can be decoupled from changes in organic matter, although responsive to plant functional type. Comparisons across sites illustrated that enzymes were most responsive in the temperate broadleaf savanna and temperate coniferous, suggesting the potential for reduced decomposition to influence changes in soil C to be higher.

### Coupling between enzyme activities and soil respiration

To test whether extracellular enzyme activities were related to soil C losses via decomposition, we compared averaged enzyme activity levels that were weighted based on tree cover at the replicate plot level with *in situ* soil respiration measurements in the temperate broadleaf savanna (n=6). We found two lines of evidence that enzyme activities were related to soil C decomposition: (i) respiration rates (scaled by temperature) were significantly lower in the burned plots than unburned plots (F_1,4_=13.9, p=0.02), and (ii) total hydrolytic C-acquisition enzyme activity was positively correlated with respiration rates (F_1,4_=7.9, p=0.048). Oxidative enzymes did not display a significant correlation (p>0.5), however. Consequently, independent measurements of soil C losses support our inference that reduction in enzyme activity approximated decomposition, at one site.

### Nitrogen limitation of enzyme activity

We next evaluated how losses of soil N could lead to lower enzymatic activity away from tree canopies and in burned plots. We tested whether (i) fire-driven changes in soil N, both total and inorganic, were positively correlated with C-acquisition activity, (ii) N-acquisition activity was higher in burned environments, and (iii) the ratio between N- and C-acquisition enzymes was higher in burned environments.

Across all sites, soil N was positively related to the total activity of hydrolytic C-acquisition enzymes (F_1,109.1_=8.8, p=0.004), but not oxidative enzymes (F_1,95.3_=1.24, p=0.27). Furthermore, IN was not as strong of a predictor as total N, but was positively correlated with hydrolytic C-acquisition enzymes in the temperate broadleaf savanna and temperate coniferous forest, but not with oxidative enzymes in any of the sites (Table S5); there was a negative effect of fire on total IN when all sites were considered (p<0.0001) (Table S6). Consequently, N in general scales positively with potential turnover of labile C compounds.

We next analyzed whether fire altered the activity of the enzyme, NAG, which can be used to acquire N in addition to C. If microbes were compensating for lower soil N, we would expect elevated activities in burned plots and/or open areas and for NAG to be negatively correlated with soil N content. In contrast to this expectation, we found that NAG was higher in the unburned plots and underneath trees (Tables 1,2, Figure 6). In all sites, NAG activity was lower in burned plots, with combustive effects of fire in the temperate coniferous savanna, temperate coniferous forest, and tropical savanna, and effects due to shifts in tree cover in the temperate broadleaf savanna (Tables 1,2, Figure 6). In unburned plots relative to burned plots, NAG activity was 941% higher in the temperate coniferous savanna, 192% higher in the temperate coniferous forest, and 101% higher in the tropical savanna (although this effect was only marginally significant at p=0.1 in the tropical savanna). Trees also tended to have elevated NAG activity in the temperate broadleaf savanna (+138%) and in the temperate coniferous forest (+182%, but only in the unburned plots). Changes due to shifting tree cover in the temperate broadleaf savanna resulted in unburned plots having 42% higher NAG activity at the landscape scale. The consistent response of NAG to fire across sites illustrates the pervasive effect of burning on the N cycle.

**Figure 6:**
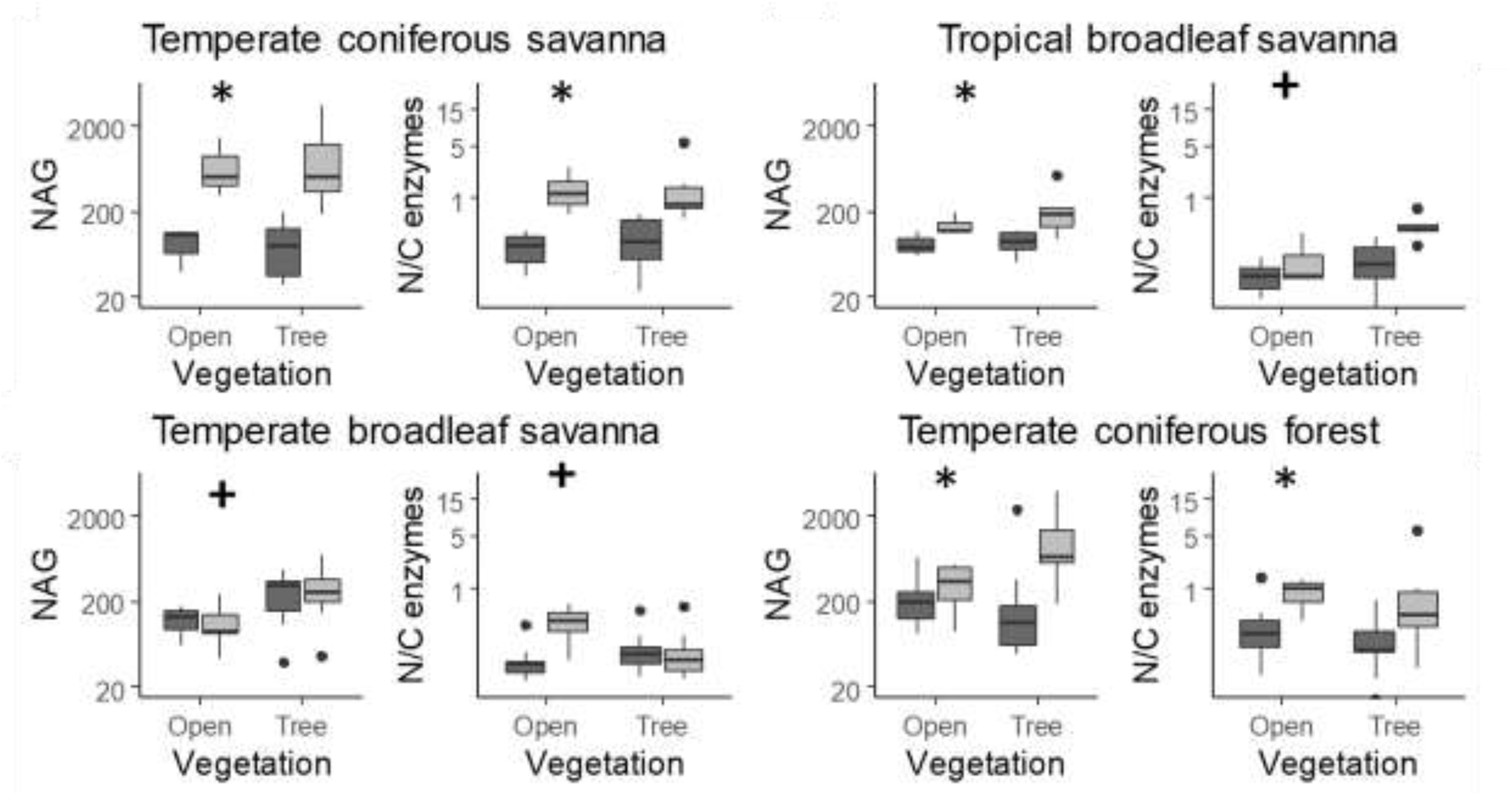
Differential fire and vegetation effects on enzyme involved in N release. Box and whiskers plots of extracellular enzyme activity of N-acetyl-β-D-glucosaminidase (NAG, degrades chitin) and its activity relative to the sum of the oxidative C acquisition enzymes (N/C enzymes, divided by 10,000 for display). Values partitioned into different fire treatments (dark grey=burned, light grey=unburned) from either underneath trees or away from tree canopies. Significant effect of fire denoted by the asterisk, significant effect of vegetation denoted by the plus symbol. Units for NAG are nmoles h^-1^ g^-1^, the ratio is unitless.

Positive correlations between NAG and soil N illustrated that N acquisition activity increased with soil N. For one, bulk soil N was positively correlated with NAG when all sites were considered (model included vegetation type and fire effects; F_1,110.6_=5.9, p=0.017), but within sites there were only correlations in the temperate broadleaf savanna and temperate coniferous forest (Table S5). There was no evidence for a saturating relationship between inorganic N and NAG activity, which would be expected if N availability was equal to organismal demand, with all four sites having positive NAG ∼ IN correlations (Table S5).

Taken together, lower soil N is coupled with reduced N acquisition and C acquisition of labile organic matter. However, there was variability in the strength of the coupling across sites, with the temperate broadleaf savanna and temperate coniferous forest tending to have a stronger coupling between bulk pools and enzyme activities.

Our final test of whether N potentially limited decomposition used the ratios between NAG vs. C-acquisition enzymes (NAG:C_hydro_ and NAG:C_oxid_). Unburned plots tended to have higher NAG activity relative to C-acquisition enzymes in the temperate broadleaf savanna and temperate coniferous forest. In both sites, trees tended to have a higher NAG:C_hydro_ and NAG:C_oxi_ than open areas, and in the temperate coniferous forest, unburned areas also had higher ratios (Tables 1,2, Figure 6). Consequently, although both C acquisition and NAG were elevated in unburned plots, fire protection increased NAG to a greater degree. In contrast, the ratios of NAG:C_hydro_ in the temperate coniferous and tropical savannas were unchanged by either fire or vegetation type, but the ratios of NAG:C_oxi_ were higher in the unburned plots in the coniferous savanna and underneath trees in the tropical savanna, largely due to higher NAG (Tables 1,2, Figure 6). The results illustrate that microbial acquisition and turnover of N is higher in unburned plots or underneath trees, potentially helping to alleviate N limitation and allowing for high C turnover.

## Discussion

Our results demonstrate that repeated burning can decrease soil C and N through reduced tree biomass inputs, and that declines in soil C and N occurred even though fire reduced the decomposition of soil organic matter. In the savannas containing C_4_ grasses, ^13^C was negatively correlated with bulk soil C, indicating lower tree inputs to soil C corresponded with lower total soil C. Along these lines, the lack of change in soil C and N in the temperate coniferous forest, where the absence of C_4_ grasses prohibited the isotopic technique, was consistent with the minimal changes in tree biomass. Moreover, in the tropical and temperate broadleaf savannas, where soils underneath trees had higher C and N than open areas, fire-driven losses of tree cover were important for fire effects on soil C and N at the landscape scale.

Our results also suggest that fire-driven suppression of extracellular enzyme activity may dampen the losses of soil C. We found a ubiquitous negative effect of fire on the activity of hydrolytic enzymes that decompose carbon compounds. Furthermore, soil CO_2_ respiration fluxes were significantly lower in burned plots and were significantly related to the activity of hydrolytic enzymes in the temperate broadleaf savanna.

Finally, fire-driven losses of N and lower N turnover suggest N limitation may be one factor contributing to reduced C turnover in burned plots. We observed (i) a significant negative effect of fire on soil N in three of the four sites (either through direct combustion effects or through fire-driven losses of tree cover), (ii) a fire-driven decline in the activity of N-acquisition enzymes in all the sites, and (iii) a positive correlation between soil N and C- and N-acquisition enzyme activity.

In total, our results indicate that fire exerts a top-down control on ecosystem biogeochemistry by reducing tree biomass and cover, but that there are also important bottom-up processes as fire can also reduce the potential turnover of C and N; consequently, the losses of soil C may partially be reduced through suppressed decomposition.

### Combustion and changing tree cover both regulate belowground changes

The role of changes in tree cover in regulating changes in soil C and N differed across the three savannas. Soil C, N, and enzymes were responsive to tree cover in the temperate and tropical broadleaf savannas, while soil chemistry in the coniferous savanna changed because of combustive effects alone. In the two broadleaf savannas, where tree cover had a strong enrichment effect, fire reduced tree abundance and canopy cover, resulting in soil C and N being lower in burned plots. The large declines in tree cover with frequent burning that tends to occur in savannas (Peterson and Reich 2001, Higgins et al. 2007, Veenendaal et al. 2018) may be one factor leading to the strong losses of soil C and N observed in a recent global meta-analysis (Pellegrini et al. 2018).

In the temperate broadleaf savanna, the lack of a fire effect on bulk soil C and N pools underneath either grasses or trees suggests that vegetation can compensate for fire-driven losses, which may be due to several factors. For one, frequently burned plots in this savanna have fire-adapted herbaceous species, shrubs, and trees (Cavender-Bares and Reich 2012), and the trees that occur in the burned plots are large mature oaks that likely have persisted since the onset of fire treatments (Peterson and Reich 2001, Peterson et al. 2007). Furthermore, the burned plots can have higher fine root biomass than the fire-protected plots, which tend to have higher amounts of coarse root biomass (*unpublished data*), potentially sustaining belowground inputs.

Resilience of vegetation to fire was most evident in the coniferous forest, where fire had little effect on aboveground pools, suggesting that there may be sustained inputs into soils. Furthermore, although fire reduced the duff layer, it remained a substantial pool (2.8 Mg ha^-1^, Figure 3), likely slowing the potential declines in inputs from biological decomposition. In contrast to no change in total C and N pools, fire reduced the activity of enzymes, potentially because of direct heating effects on microbial biomass.

The results from the temperate coniferous forest contrast with a recent meta-analysis that found that fire enriches soil C and N in needleleaf ecosystems (Pellegrini et al. 2018). The post-fire leaching of C and N that has been observed in our site (Chorover et al. 1994, Engle et al. 2008) may play a role in limiting the efficiency of redistribution from ash into soil. Furthermore, the lack of a fire effect on total soil N is consistent with results from a longer-term study on natural variability in fire frequency in ponderosa pine/Douglas-fir forests (DeLuca and Sala 2006). Our results may contrast with those in the global meta-analysis because the majority of the needleleaf sites in the meta-analysis were from longleaf and loblolly pine woodlands in the southeastern United States.

### Changes in extracellular enzymes

In all sites, fire reduced potential extracellular enzyme activity, which may reduce further losses of soil C. First, when C enzymes were analyzed individually, fire and/or the absence of trees reduced the activity of at least one enzyme in all sites, although the strength of the responses tended to vary across sites. Second, in all comparisons, fire never increased the activity of enzymes, contrasting with previous studies (Boerner et al. 2000, 2006, Rietl and Jackson 2012). Third, respiration rates in the temperate broadleaf savanna also declined with fire and were correlated with hydrolytic enzyme activity across the replicate plots. Consequently, our results agree with some previous studies that have found enzyme activity is reduced in some cases by single fires (Waldrop and Harden 2008) as well as more consistently by repeated burning (Eivazi and Bayan 1996, Ajwa et al. 1999, Boerner et al. 2005). Consequently, negative fire effects on enzymes may be a general trend.

We found stark differences in the responses of the enzymes involved in processing labile C compounds of cellulose, hemicellulose, and starch vs. recalcitrant C compounds of lignin and polyphenols (Sinsabaugh et al. 2008). Total oxidative enzyme activity never declined due to fire alone. However, there was higher oxidative activity underneath trees in the temperate broadleaf savanna and coniferous forest. The link between tree cover and oxidative enzyme activity may explain the declines in oxidative activity after burning at the landscape scale in other studies (Waldrop and Harden 2008, Holden et al. 2013).

The losses of N and reduction in N turnover caused by fire and reduced tree cover may contribute to reduced C turnover. N likely limits C turnover given that lower soil N was associated with lower hydrolytic C-acquisition enzyme activities even when controlling for direct fire and/or vegetation effects. Moreover, fire likely amplifies low N conditions by reducing NAG activity in all sites either due to direct combustive effects or fire-driven shifts in tree cover. Finally, potential N turnover was more sensitive to fire and vegetation type than potential C turnover, illustrated by the higher C_hydro_:NAG ratios in burned plots or away from tree canopies. Consequently, fire disproportionately suppressed N turnover and may promote N limitation.

It is unlikely that the observed declines in enzyme activities are artifacts of sampling times since fire, given that the largest effects of fire on enzyme activities occurred in the two sites with the longest time since fire (5-6 years in the temperate coniferous forest and coniferous savanna). Consequently, the recovery of enzymes in a few years after a single fire (e.g., (Gutknecht et al. 2010)), does not always translate into long-term recovery when burning is repeated and suggests fire leaves a legacy effect on microbial processes.

## Conclusions

Even though fire consistently combusts plant biomass and volatilizes C and N to the atmosphere across ecosystems, repeated burning does not always lead to losses of soil C and N (Pellegrini et al. 2018). Our analyses support the hypothesis that consumption of aboveground biomass is a key factor regulating fire’s effect on soils (McKee 1982, Guinto et al. 2001, Reich et al. 2001, Holdo et al. 2009b, Pellegrini et al. 2014), with inter-site differences in fire-driven declines in tree biomass explaining variability in the effect of fire on soil C and N content. However, bottom-up factors can also be important, given that the potential activities of enzymes that break down organic matter were lower in burned plots and away from trees, potentially reducing soil C losses and depending on low soil N. However, the higher potential decomposition activity in unburned plots and underneath trees does not appear to suppress soil C accumulation from increased biomass inputs, illustrating the complex interactions between above- and belowground processes in determining the net-changes of soil C.

## Acknowledgements

A. P. was supported by a NOAA Climate and Global Change postdoctoral fellowship program and the USDA National Institute of Food and Agriculture postdoctoral fellowship program. R. J. received support from the Gordon and Betty Moore Foundation. The experiments in the sites were organized and funded through the Cedar Creek Long Term Ecological Research program, the National Park Service, and South African National Parks. We thank Bryce Currey, Katerina Georgiou, Natalie Gross, Lars Hedin, Devin McMahon, and Mingzhen Lu for assistance in the field.

## Appendix: Tables S1-S6

**Table S1:** Means and standard errors of total %C, %N, C:N, and ^13^C across the sites, fire treatments, and vegetation types. “T.” or “Tr.” indicate temperate or tropical, respectively; “conif.” vs. “broad.” indicate coniferous vs. broadleaf tree types, respectively; “sav” vs. “for” indicate savanna vs. forest, respectively.

| *Site* | *Fire* | *Vegetation* | *%C* | *SE C* | *%N* | *SE N* | *C:N* | *SE C:N* | *$^{13}\text{C}$* | *SE $^{13}\text{C}$* |
| --- | --- | --- | --- | --- | --- | --- | --- | --- | --- | --- |
| *T. conif. sav* | High | Open | 1.65 | 0.48 | 0.093 | 0.018 | 17.09 | 3.43 | -17.68 | 2.33 |
| *T. conif. sav* | High | Tree | 2.22 | 0.52 | 0.085 | 0.010 | 27.24 | 3.20 | -16.06 | 1.37 |
| *T. conif. sav* | Low | Open | 3.68 | 0.36 | 0.267 | 0.028 | 13.89 | 0.73 | -25.99 | 0.32 |
| *T. conif. sav* | Low | Tree | 4.68 | 0.59 | 0.240 | 0.013 | 19.56 | 2.35 | -24.44 | 0.48 |
| *T. broad. sav* | High | Open | 1.83 | 0.66 | 0.139 | 0.040 | 12.79 | 1.01 | -22.54 | 0.80 |
| *T. broad. sav* | High | Tree | 2.77 | 0.94 | 0.201 | 0.059 | 13.52 | 0.61 | -25.82 | 0.86 |
| *T. broad. sav* | Low | Open | 1.18 | 0.26 | 0.113 | 0.015 | 9.87 | 0.80 | -23.60 | 1.18 |
| *T. broad. sav* | Low | Tree | 3.22 | 0.71 | 0.227 | 0.043 | 13.60 | 0.76 | -27.08 | 0.24 |
| *Tr. broad. sav* | High | Open | 0.80 | 0.11 | 0.063 | 0.009 | 12.57 | 0.56 | -16.43 | 0.56 |
| *Tr. broad. sav* | High | Tree | 1.46 | 0.12 | 0.099 | 0.006 | 14.58 | 0.16 | -20.85 | 0.20 |
| *Tr. broad. sav* | Low | Open | 1.04 | 0.10 | 0.087 | 0.007 | 12.01 | 0.76 | -18.41 | 0.42 |
| *Tr. broad. sav* | Low | Tree | 1.88 | 0.08 | 0.139 | 0.006 | 13.49 | 0.68 | -21.99 | 0.35 |
| *T. conif. for* | High | Open | 4.77 | 0.62 | 0.251 | 0.032 | 18.98 | 0.10 | -24.14 | 0.09 |
| *T. conif. for* | High | Tree | 3.54 | 0.33 | 0.200 | 0.018 | 17.29 | 0.23 | -23.89 | 0.26 |
| *T. conif. for* | Low | Open | 5.29 | 1.50 | 0.368 | 0.084 | 16.87 | 5.96 | -25.03 | 0.06 |
| *T. conif. for* | Low | Tree | 7.36 | 2.63 | 0.315 | 0.099 | 22.68 | 0.73 | -24.35 | 0.03 |

**Table S2:** Changes in woody vegetation structure and its impact on soil C and N pools at the landscape scale. Means and standard errors for basal area, density of individuals, and tree cover, which is expressed as a percentage of landscape covered by tree canopies. The %Δ C and %Δ N represent the impact of considering tree-cover weighted C and N calculations on estimates of bulk soil concentrations at the landscape scale compared to an unweighted average across grass and tree soil samples.

| ***Tropical broadleaf savanna*** | | *BA*<br>($m^2 ha^{-1}$) | *Indiv*<br>($\# ha^{-1}$) | *TC*<br>(%) | %Δ C | %Δ N |
| --- | --- | --- | --- | --- | --- | --- |
| *High* | Means | 0.08 | 276 | 1.3 | -28.4 | -21.6 |
| *High* | SEMs | 0.03 | 32 | 0.5 | 7.8 | 7.1 |
| *Low* | Means | 4.94 | 1079 | 78.1 | 16.5 | 13.2 |
| *Low* | SEMs | 0.48 | 77 | 7.7 | 5.9 | 4.0 |
| ***Temperate broadleaf savanna*** |  |  |  |  |  |  |
| *High* | Means | 9.20 | 118 | 45.6 | -1.2 | -0.7 |
| *High* | SEMs | 3.34 | 33 | 5.2 | 3.2 | 3.1 |
| *Low* | Means | 22.53 | 694 | 83.8 | 30.7 | 22.4 |
| *Low* | SEMs | 3.65 | 176 | 5.4 | 11.4 | 9.2 |

**Table S3:** Statistical analyses testing the effect of fire on tree cover, density, and basal area using ANOVAs. ANOVAs run on replicate plot means and analyzed within each site.

| <u>*Tropical broadleaf savanna*</u> | <u>*Fire effects*</u> |
| --- | --- |
| *Individuals (# ha<sup>-1</sup>)* | F<sub>1,4</sub>=73.7, p=0.001 |
| *Tree cover %* | F<sub>1,4</sub>=55.9, p=0.002 |
| *Basal area (m<sup>2</sup> ha<sup>-1</sup>)* | F<sub>1,4</sub>=56.1, p=0.002 |
| <u>*Temperate broadleaf savanna*</u> |  |
| *Individuals (# ha<sup>-1</sup>)* | F<sub>1,5</sub>=16.1, p=0.010 |
| *Tree cover %* | F<sub>1,5</sub>=16.1, p=0.010 |
| *Basal area (m<sup>2</sup> ha<sup>-1</sup>)* | F<sub>1,5</sub>=4.66, p=0.083 |

**Table S4:**
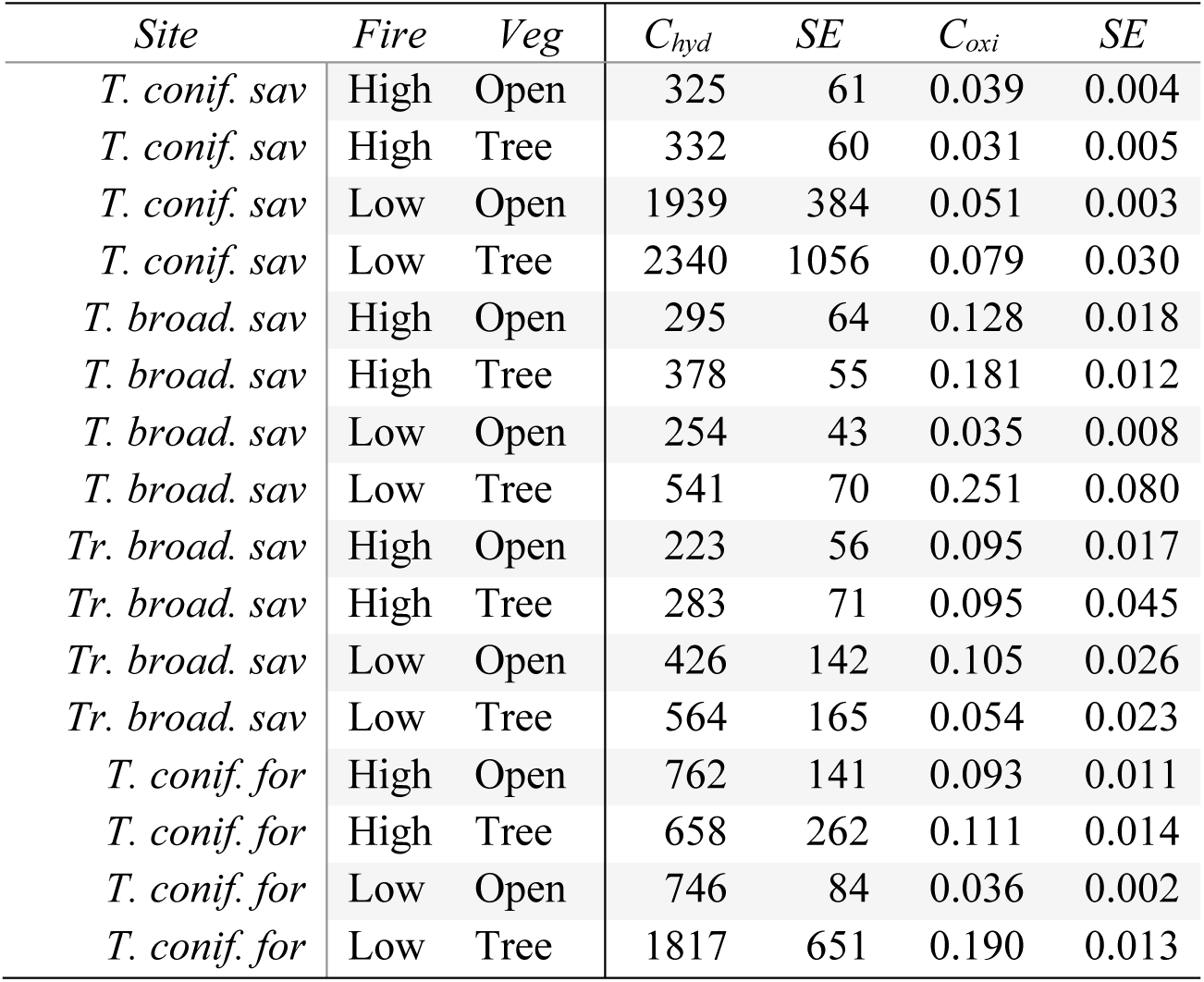
Means and standard errors for enzymes involved in carbon acquisition partitioned into hydrolytic (C_hyd_) and oxidative (C_oxi_) C acquisition enzymes. “T.” or “Tr.” indicate temperate or tropical, respectively; “conif.” vs. “broad.” indicate coniferous vs. broadleaf tree types, respectively; “sav” vs. “for” indicate savanna vs. forest, respectively.

| Site | Fire | Veg | $C_{hyd}$ | SE | $C_{oxi}$ | SE |
| --- | --- | --- | --- | --- | --- | --- |
| *T. conif. sav* | High | Open | 325 | 61 | 0.039 | 0.004 |
| *T. conif. sav* | High | Tree | 332 | 60 | 0.031 | 0.005 |
| *T. conif. sav* | Low | Open | 1939 | 384 | 0.051 | 0.003 |
| *T. conif. sav* | Low | Tree | 2340 | 1056 | 0.079 | 0.030 |
| *T. broad. sav* | High | Open | 295 | 64 | 0.128 | 0.018 |
| *T. broad. sav* | High | Tree | 378 | 55 | 0.181 | 0.012 |
| *T. broad. sav* | Low | Open | 254 | 43 | 0.035 | 0.008 |
| *T. broad. sav* | Low | Tree | 541 | 70 | 0.251 | 0.080 |
| *Tr. broad. sav* | High | Open | 223 | 56 | 0.095 | 0.017 |
| *Tr. broad. sav* | High | Tree | 283 | 71 | 0.095 | 0.045 |
| *Tr. broad. sav* | Low | Open | 426 | 142 | 0.105 | 0.026 |
| *Tr. broad. sav* | Low | Tree | 564 | 165 | 0.054 | 0.023 |
| *T. conif. for* | High | Open | 762 | 141 | 0.093 | 0.011 |
| *T. conif. for* | High | Tree | 658 | 262 | 0.111 | 0.014 |
| *T. conif. for* | Low | Open | 746 | 84 | 0.036 | 0.002 |
| *T. conif. for* | Low | Tree | 1817 | 651 | 0.190 | 0.013 |

**Table S4:** Means and standard errors of NAG and its activity relative to C acquisition enzymes (C_hyd_: hydrolytic enzymes; C_oxi_: oxidative enzymes; the oxidative ratio is *10000 for display purposes).

| Site | Fire | Veg | NAG | SE | $C_{hyd}:NAG$ | SE | $C_{oxi}:NAG$ | SE |
| --- | --- | --- | --- | --- | --- | --- | --- | --- |
| *T. conif. sav* | High | Open | 88 | 25 | 4.03 | 0.64 | 5.9 | 2.6 |
| *T. conif. sav* | High | Tree | 93 | 24 | 4.42 | 1.44 | 5.8 | 2.0 |
| *T. conif. sav* | Low | Open | 773 | 360 | 3.26 | 0.93 | 1.0 | 0.4 |
| *T. conif. sav* | Low | Tree | 1122 | 720 | 2.93 | 0.54 | 1.1 | 0.1 |
| *T. broad. sav* | High | Open | 126 | 10 | 2.46 | 0.45 | 10.9 | 1.4 |
| *T. broad. sav* | High | Tree | 274 | 65 | 1.76 | 0.36 | 7.9 | 2.0 |
| *T. broad. sav* | Low | Open | 119 | 17 | 2.33 | 0.38 | 3.4 | 1.3 |
| *T. broad. sav* | Low | Tree | 305 | 41 | 1.87 | 0.10 | 9.4 | 1.9 |
| *Tr. broad. sav* | High | Open | 86 | 18 | 2.54 | 0.12 | 12.7 | 4.4 |
| *Tr. broad. sav* | High | Tree | 91 | 14 | 3.13 | 0.32 | 10.8 | 5.3 |
| *Tr. broad. sav* | Low | Open | 144 | 25 | 2.81 | 0.44 | 8.3 | 2.7 |
| *Tr. broad. sav* | Low | Tree | 231 | 102 | 2.71 | 0.59 | 2.6 | 0.2 |
| *T. conif. for* | High | Open | 238 | 61 | 3.62 | 0.50 | 5.3 | 2.2 |
| *T. conif. for* | High | Tree | 334 | 190 | 2.94 | 0.34 | 7.9 | 3.1 |
| *T. conif. for* | Low | Open | 348 | 37 | 2.10 | 0.19 | 1.3 | 0.2 |
| *T. conif. for* | Low | Tree | 1320 | 557 | 1.97 | 0.63 | 2.8 | 1.3 |

**Table S5:** Analyses of mixed effects regressions between enzyme activity and soil carbon (C), nitrogen (N), and inorganic N (IN) with statistical significance determined by Satterthwaite’s method. Each relationship was determined using model selection including the potential effects of fire treatment and vegetation type on enzyme activity and to modify the regression. First degrees of freedom always equals one. An asterisk indicates that the regression depended on an interaction with fire treatment. “T.” or “Tr.” indicate temperate or tropical, respectively; “conif.” vs. “broad.” indicate coniferous vs. broadleaf tree types, respectively; “sav” vs. “for” indicate savanna vs. forest, respectively.

| *Hydrolytic~%C* | *df* | *F* | *p value* |
| --- | --- | --- | --- |
| *T. conif. sav* | 38.00 | 23.40 | <0.0001 |
| *T. broad. sav* | 38.00 | 13.36 | 0.001 |
| *Tr. broad. sav* | 12.95 | 16.98 | 0.001 |
| *T. conif. for* | 34.00 | 43.01 | <0.0001 |
| *Oxidative~%C* |  |  |  |
| *T. conif. sav* | 14.00 | 4.29 | 0.057 |
| *T. broad. sav* | 23.21 | 16.90 | <0.001 |
| *Tr. broad. sav* | 12.21 | 3.96 | 0.070 |
| *T. conif. for* | 37.00 | 0.57 | 0.453 |
| *Hydrolytic~%N* |  |  |  |
| *T. conif. sav* | 13 | 1.25 | 0.2833 |
| *T. broad. sav* | 38 | 12.3 | 0.001176 |
| *Tr. broad. sav* | 13 | 16.8 | 0.001246 |
| *T. conif. for* | 34.2 | 4.23 | 0.04751 |
| *Oxidative~%N* |  |  |  |
| *T. conif. sav\** | 13 | 1.57 | 0.233 |
| *T. broad. sav* | 25.7 | 4.24 | 0.049823 |
| *Tr. broad. sav* | 12.5 | 0.1 | 0.7597 |
| *T. conif. for* | 36 | 0.03 | 0.856129 |
| *NAG~%N* |  |  |  |
| *T. conif. sav* | 13.95 | 0.10 | 0.762 |
| *T. broad. sav* | 37.00 | 4.33 | 0.044 |
| *Tr. broad. sav* | 13.58 | 0.70 | 0.417 |
| *T. conif. for* | 35.99 | 25.53 | <0.0001 |
| *NAG~IN* |  |  |  |
| *T. conif. sav\** | 8.00 | 14.14 | 0.006 |
| *T. broad. sav* | 39.00 | 6.35 | 0.016 |
| *Tr. broad. sav\** | 13.16 | 3.61 | 0.080 |
| *T. conif. for* | 35.97 | 12.45 | 0.001 |

**Table S6:** means and standard errors of fire and vegetation type effects on inorganic nitrogen concentrations partitioned into nitrate (NO_3_) and ammonium (NH_4_) as well as their sum (IN).

| *Site* | *Fire* | *Veg. type* | *NO<sub>3</sub>* | *SE NO<sub>3</sub>* | *NH<sub>4</sub>* | *SE NH<sub>4</sub>* | *IN* | *SE IN* |
| --- | --- | --- | --- | --- | --- | --- | --- | --- |
| *T. conif. sav* | High | Open | 0.96 | 0.096 | 2.24 | 0.30 | 3.20 | 0.379 |
| *T. conif. sav* | High | Tree | 0.46 | 0.280 | 2.50 | 0.10 | 2.96 | 0.380 |
| *T. conif. sav* | Low | Open | 1.93 | 0.482 | 25.90 | 13.40 | 27.83 | 13.836 |
| *T. conif. sav* | Low | Tree | 0.68 | 0.125 | 7.79 | 0.59 | 8.47 | 0.709 |
| *T. broad. sav* | High | Open | 0.20 | 0.028 | 1.95 | 0.43 | 2.15 | 0.437 |
| *T. broad. sav* | High | Tree | 0.31 | 0.087 | 3.19 | 0.58 | 3.50 | 0.641 |
| *T. broad. sav* | Low | Open | 0.23 | 0.067 | 2.40 | 0.31 | 2.63 | 0.243 |
| *T. broad. sav* | Low | Tree | 0.33 | 0.133 | 3.40 | 0.06 | 3.72 | 0.174 |
| *Tr. broad. sav* | High | Open | 0.01 | 0.003 | 0.37 | 0.05 | 0.38 | 0.054 |
| *Tr. broad. sav* | High | Tree | 0.02 | 0.001 | 0.31 | 0.002 | 0.34 | 0.001 |
| *Tr. broad. sav* | Low | Open | 0.01 | 0.005 | 0.31 | 0.06 | 0.32 | 0.068 |
| *Tr. broad. sav* | Low | Tree | 0.02 | 0.002 | 0.37 | 0.01 | 0.39 | 0.016 |
| *T. conif. for* | High | Open | 2.89 | 0.577 | 6.40 | 2.15 | 9.29 | 1.580 |
| *T. conif. for* | High | Tree | 1.74 | 0.502 | 6.92 | 1.36 | 8.65 | 1.570 |
| *T. conif. for* | Low | Open | 1.82 | 0.710 | 11.78 | 2.87 | 13.60 | 2.571 |
| *T. conif. for* | Low | Tree | 1.11 | 0.595 | 19.99 | 7.52 | 21.09 | 7.605 |

## References

Ajwa, H. A., C. J. Dell, and C. W. Rice. 1999. Changes in enzyme activities and microbial biomass of tallgrass prairie soil as related to burning and nitrogen fertilization. Soil Biology and Biochemistry 31:769–777.

Andela, N., D. C. Morton, L. Giglio, Y. Chen, G. R. van der Werf, P. S. Kasibhatla, R. S. DeFries, G. J. Collatz, S. Hantson, S. Kloster, D. Bachelet, M. Forrest, G. Lasslop, F. Li, S. Mangeon, J. R. Melton, C. Yue, and J. T. Randerson. 2017. A human-driven decline in global burned area. Science 356:1356–1362.

Archibald, S., C. E. R. Lehmann, J. L. Gómez-Dans, and R. A. Bradstock. 2013. Defining pyromes and global syndromes of fire regimes. Proceedings of the National Academy of Sciences 110:6442–6447.

Bates, D., M. Mächler, B. Bolker, and S. Walker. 2014. Fitting Linear Mixed-Effects Models using lme 4.

Belsky, A. J. A. J., R. G. Amundson, J. M. Duxbury, S. J. Riha, A. R. Ali, and S. M. Mwonga. 1989. The effects of trees on their physical, chemical and biological environments in a semi-arid savanna in Kenya. Journal of Applied Ecology 26:1005–1024.

Biggs, H. C., T. T. Dunne, and N. Govender. 2003. Experimental burn plot trial in the Kruger National Park?: history, experimental design and suggestions for data analysis. Koedoe 1.

Bird, M. I., and P. L. Ascough. 2012. Isotopes in pyrogenic carbon: A review. Organic Geochemistry 42:1529–1539.

Boerner, R. E. J., J. A. Brinkman, and A. Smith. 2005. Seasonal variations in enzyme activity and organic carbon in soil of a burned and unburned hardwood forest. Soil Biology and Biochemistry 37:1419–1426.

Boerner, R. E. J., K. L. M. Decker, and E. K. Sutherland. 2000. Prescribed burning effects on soil enzyme activity in a southern Ohio hardwood forest: a landscape-scale analysis. Soil Biology and Biochemistry 32:899–908.

Boerner, R. E., T. A. Waldrop, and V. B. Shelburne. 2006. Wildfire mitigation strategies affect soil enzyme activity and soil organic carbon in loblolly pine (Pinus taeda) forests. Canadian Journal of Forest Research 36:3148–3154.

Bond-Lamberty, B., S. D. Peckham, D. E. Ahl, and S. T. Gower. 2007. Fire as the dominant driver of central Canadian boreal forest carbon balance. Nature 450:89–92.

Bowman, D. M. J. S., J. K. Balch, P. Artaxo, W. J. Bond, J. M. Carlson, M. A. Cochrane, C. M. D’Antonio, R. S. DeFries, J. C. Doyle, S. P. Harrison, F. H. Johnston, J. E. Keeley, M. A. Krawchuk, C. A. Kull, J. B. Marston, M. A. Moritz, I. C. Prentice, C. I. Roos, A. C. Scott, T. W. Swetnam, G. R. van der Werf, and S. J. Pyne. 2009. Fire in the Earth System. Science 324:481–484.

Caprio, A. C., and T. W. Swetnam. 1995. Historic fire regimes along an elevational gradient on the west slope of the Sierra Nevada, California. Pages 173–179 in J. K.. Brown, R. W. Mutch, C. W. Spoon, and R. H. Wakimoto, editors. Proceedings: symposium on fire in wilderness and park management; 1993 March 30-April 1, Missoula, MT. Gen. Tech. Rep. INT-GTR-320. Ogden.

Cavender-Bares, J., and P. B. Reich. 2012. Shocks to the system: community assembly of the oak savanna in a 40-year fire frequency experiment. Ecology 93:S52–S69.

Chorover, J., P. M. Vitousek, D. A. Everson, A. M. Esperanza, and D. Turner. 1994. Solution chemistry profiles of mixed-conifer forests before and after fire. Source: Biogeochemistry 26:115–144.

Coetsee, C., W. J. Bond, and E. C. February. 2010. Frequent fire affects soil nitrogen and carbon in an African savanna by changing woody cover. Oecologia 162:1027–34.

DeLuca, T. H., and A. Sala. 2006. Frequent fire alters nitrogen transformations in ponderosa pine stands of the inland northwest. Ecology 87:2511–2522.

Dennison, P. E., S. C. Brewer, J. D. Arnold, and M. A. Moritz. 2014. Large wildfire trends in the western United States, 1984–2011. Geophysical Research Letters 41:2928–2933.

Dijkstra, F. A., K. Wrage, S. E. Hobbie, and P. B. Reich. 2006. Tree patches show greater N losses but maintain higher soil N availability than grassland patches in a frequently burned oak savanna. Ecosystems 9:441–452.

Eivazi, F., and M. R. Bayan. 1996. Effects of long-term prescribed burning on the activity of select soil enzymes in an oak-hickory forest. Canadian Journal of Forest Research 26:1799–1804.

Engle, D. L., J. O. Sickman, C. M. Moore, A. M. Esperanza, J. M. Melack, and J. E. Keeley. 2008. Biogeochemical legacy of prescribed fire in a giant sequoia - Mixed conifer forest: A 16-year record of watershed balances. Journal of Geophysical Research: Biogeosciences.

Guinto, D. F., Z. H. Xu, P. G. Saffigna, A. P. N. House, P. G. Saffigna, and A. P. N. House. 2001. Soil chemical properties and forest floor nutrients under repeated prescribed-burning in eucalypt forests of south-east Queensland, Australia. New Zealand Journal of Forestry Science 31:170–187.

Gutknecht, J. L. M., H. A. L. Henry, and T. C. Balser. 2010. Inter-annual variation in soil extra-cellular enzyme activity in response to simulated global change and fire disturbance. Pedobiologia 53:283–293.

Hart, S. C., A. T. Classen, and R. J. Wright. 2005. Long-term interval burning alters fine root and mycorrhizal dynamics in a ponderosa pine forest. Journal of Applied Ecology 42:752–761.

Higgins, S. I., W. J. Bond, E. C. February, A. Bronn, D. I. W. Euston-Brown, B. Enslin, N. Govender, L. Rademan, S. O’Regan, and A. L. F. Potgieter. 2007. Effects of four decades of fire manipulation on woody vegetation structure in savanna. Ecology 88:1119–1125.

Hobbie, S. E., W. C. Eddy, C. R. Buyarski, E. C. Adair, M. L. Ogdahl, and P. Weisenhorn. 2012. Response of decomposing litter and its microbial community to multiple forms of nitrogen enrichment. Ecological Monographs 82:389–405.

Holden, S. R., A. Gutierrez, and K. K. Treseder. 2013. Changes in Soil Fungal Communities, Extracellular Enzyme Activities, and Litter Decomposition Across a Fire Chronosequence in Alaskan Boreal Forests. Ecosystems 16:34–46.

Holdo, R. M., R. D. Holt, and J. M. Fryxell. 2009a. Grazers, browsers, and fire influence the extent and spatial pattern of tree cover in the Serengeti. Ecological Applications 19:95–109.

Holdo, R. M., M. C. Mack, and S. G. Arnold. 2011. Tree canopies explain fire effects on soil nitrogen, phosphorus and carbon in a savanna ecosystem. Journal of Vegetation Science.

Holdo, R. M., A. R. E. Sinclair, A. P. Dobson, K. L. Metzger, B. M. Bolker, M. E. Ritchie, and R. D. Holt. 2009b. A disease-mediated trophic cascade in the Serengeti and its implications for ecosystem C. PLoS Biol 7:e1000210.

Jackson, R. B., K Lajtha, S. E. Crow, G. Hugelius, M. G. Kramer, G. Piñeiro. The ecology of soil carbon: pools, vulnerabilities, and biotic and abiotic controls. Annual Review of Ecology, Evolution, and Systematics 48:419–445.

Kauffman, J. B., D. L. Cummings, D. E. Ward, and R. Babbitt. 1995. Fire in the Brazilian Amazon: 1. Biomass, nutrient pools, and losses in slashed primary forests. Oecologia 104:397–408.

Kowaljow, E., M. S. Morales, J. I. Whitworth-Hulse, S. R. Zeballos, M. A. Giorgis, M. Rodríguez Catón, and D. E. Gurvich. 2018. A 55-year-old natural experiment gives evidence of the effects of changes in fire frequency on ecosystem properties in a seasonal Subtropical Dry forest. Land Degradation & Development.

Kuznetsova, A., P. B. Brockhoff, and R. H. B. Christensen. 2017. lmerTest Package: Tests in Linear Mixed Effects Models. Journal of Statistical Software 82:1548–7660.

Ludwig, F., H. de Kroon, F. Berendse, and H. H. T. Prins. 2004. The influence of savanna trees on nutrient, water and light availability and the understorey vegetation. Plant Ecology (formerly Vegetatio) 170:93–105.

van Mantgem, P. J., A. C. Caprio, N. L. Stephenson, and A. J. Das. 2016. Does prescribed fire promote resistance to drought in low elevation forests of the Sierra Nevada, California, USA? Fire Ecology 12:13–25.

McKee, W. H. 1982. Changes in soil fertility following prescribed burning on coastal plain pine sites. Southeastern Forest Experiment Station, Asheville, NC.

Melillo, J. M., J. D. Aber, and J. F. Muratore. 1982. Nitrogen and Lignin Control of Hardwood Leaf Litter Decomposition Dynamics. Ecology 63:621–626.

Muqaddas, B., X. Zhou, T. Lewis, C. Wild, and C. Chen. 2015. Long-term frequent prescribed fire decreases surface soil carbon and nitrogen pools in a wet sclerophyll forest of Southeast Queensland, Australia. Science of The Total Environment 536:39–47.

Neff, J., J. Harden, and G. Gleixner. 2005. Fire effects on soil organic matter content, composition, and nutrients in boreal interior Alaska. Canadian Journal of Forest 35:2178–2187.

Ojima, D. S., D. S. Schimel, W. J. Parton, and C. E. Owensby. 1994. Long- and short-term effects of fire on nitrogen cycling in tallgrass prairie. Biogeochemistry 24:67–84.

Pellegrini, A. F. A. A. F. A., W. A. W. A. Hoffmann, and A. C. A. C. Franco. 2014. Carbon accumulation and nitrogen pool recovery during transitions from savanna to forest in central Brazil. Ecology 95:342–352.

Pellegrini, A. F. A., A. Ahlström, S. E. Hobbie, P. B. Reich, L. P. Nieradzik, A. C. Staver, B. C. Scharenbroch, A. Jumpponen, W. R. L. Anderegg, J. T. Randerson, and R. B. Jackson. 2018. Fire frequency drives decadal changes in soil carbon and nitrogen and ecosystem productivity. Nature 553:194–198.

Peterson, D. W., and P. B. Reich. 2001. Prescribed fire in oak savanna: fire frequency effects on stand structure and dynamics. Ecological Applications 11:914–927.

Peterson, D. W., P. B. Reich, K. J. Wrage, and J. Franklin. 2007. Plant functional group responses to fire frequency and tree canopy cover gradients in oak savannas and woodlands. Journal of Vegetation Science 18:3–12.

Reich, P. B., D. W. Peterson, D. A. Wedin, and K. Wrage. 2001. Fire and vegetation effects on productivity and nitrogen cycling across a forest-grassland continuum. Ecology 82:1703–1719.

Rietl, A. J., and C. R. Jackson. 2012. Effects of the ecological restoration practices of prescribed burning and mechanical thinning on soil microbial enzyme activities and leaf litter decomposition. Soil Biology and Biochemistry 50:47–57.

Saiya-Cork, K., R. Sinsabaugh, and D. Zak. 2002. The effects of long term nitrogen deposition on extracellular enzyme activity in an Acer saccharum forest soil. Soil Biology and Biochemistry 34:1309–1315.

Schwilk, D. W., and A. C. Caprio. 2011. Scaling from leaf traits to fire behaviour: community composition predicts fire severity in a temperate forest. Journal of Ecology 99:970–980.

Sinsabaugh, R. L., R. K. Antibus, A. E. Linkins, C. A. McClaugherty, L. Rayburn, D. Repert, and T. Weiland. 1992. Wood decomposition over a first-order watershed: Mass loss as a function of lignocellulase activity. Soil Biology and Biochemistry 24:743–749.

Sinsabaugh, R. L., and J. J. Follstad Shah. 2011. Ecoenzymatic stoichiometry of recalcitrant organic matter decomposition: the growth rate hypothesis in reverse. Biogeochemistry 102:31–43.

Sinsabaugh, R. L., C. L. Lauber, M. N. Weintraub, B. Ahmed, S. D. Allison, C. Crenshaw, A. R. Contosta, D. Cusack, S. Frey, M. E. Gallo, T. B. Gartner, S. E. Hobbie, K. Holland, B. L. Keeler, J. S. Powers, M. Stursova, C. Takacs-Vesbach, M. P. Waldrop, M. D. Wallenstein, D. R. Zak, and L. H. Zeglin. 2008. Stoichiometry of soil enzyme activity at global scale. Ecology letters 11:1252–64.

Stephenson, N. L. 1988. Climatic control of vegetation distribution: the role of the water balance with examples from North America and Sequoia National Park, California. Cornell University.

Swetnam, T. W., C. H. Baisan, A. C. Caprio, P. M. Brown, R. Touchan, R. Scott Anderson, and D. J. Hallett. 2009. Multi-millennial fire history of the giant forest, Sequoia National Park, California, USA. Fire Ecology.

Treseder, K. K., M. C. Mack, and A. Cross. 2004. Relationships among fires, fungi, and soil dynamics in Alaskan boreal forests. Ecological Applications 14:1826–1838.

Turner, M. G., E. A. H. Smithwick, K. L. Metzger, D. B. Tinker, and W. H. Romme. 2007. Inorganic nitrogen availability after severe stand-replacing fire in the Greater Yellowstone ecosystem. Proceedings of the National Academy of Sciences 104:4782–4789.

USDI National Park Service. 2003. Fire Monitoring Handbook. Boise (ID): Fire Management Program Center, National Interagency Fire Center. 274p.

Veenendaal, E. M., M. Torello-Raventos, H. S. Miranda, N. M. Sato, I. Oliveras, F. van Langevelde, G. P. Asner, and J. Lloyd. 2018. On the relationship between fire regime and vegetation structure in the tropics. New Phytologist 218:153–166.

Waldrop, M. P., and J. W. Harden. 2008. Interactive effects of wildfire and permafrost on microbial communities and soil processes in an Alaskan black spruce forest. Global Change Biology 14:2591–2602.

van der Werf, G. R., J. T. Randerson, L. Giglio, T. T. van Leeuwen, Y. Chen, B. M. Rogers, M. Mu, M. J. E. van Marle, D. C. Morton, G. J. Collatz, R. J. Yokelson, and P. S. Kasibhatla. 2017. Global fire emissions estimates during 1997-2016. Earth System Science Data in press.

Westerling, A. L., H. G. Hidalgo, D. R. Cayan, and T. W. Swetnam. 2006. Warming and earlier spring increase western US forest wildfire activity. Science 313:940–943.

